# Maximum Likelihood Inference of Time-scaled Cell Lineage Trees with Mixed-type Missing Data

**DOI:** 10.1101/2024.03.05.583638

**Authors:** Uyen Mai, Gillian Chu, Benjamin J. Raphael

**Affiliations:** Department of Computer Science, Princeton University, Princeton, NJ 08544, USA

## Abstract

Recent dynamic lineage tracing technologies combine CRISPR-based genome editing with single-cell sequencing to track cell divisions during development. A key computational problem in dynamic lineage tracing is to infer a cell lineage tree from the measured CRISPR-induced mutations. Three features of dynamic lineage tracing data distinguish this problem from standard phylogenetic tree inference. First, the CRISPR-editing process modifies a genomic location exactly once. This *non-modifiable* property is not well described by the time-reversible models commonly used in phylogenetics. Second, as a consequence of non-modifiability, the number of mutations per time unit decreases over time. Third, CRISPR-based genome-editing and single-cell sequencing results in high rates of both heritable and non-heritable (dropout) missing data. To model these features, we introduce the Probabilistic Mixed-type Missing (PMM) model. We describe an algorithm, LAML (Lineage Analysis via Maximum Likelihood), to search for the maximum likelihood (ML) tree under the PMM model. LAML combines an Expectation Maximization (EM) algorithm with a heuristic tree search to jointly estimate tree topology, branch lengths and missing data parameters. We derive a closed-form solution for the M-step in the case of no heritable missing data, and a block coordinate ascent approach in the general case which is more efficient than the standard General Time Reversible (GTR) phylogenetic model. On simulated data, LAML infers more accurate tree topologies and branch lengths than existing methods, with greater advantages on datasets with higher ratios of heritable to non-heritable missing data. We show that LAML provides unbiased *time-scaled* estimates of branch lengths. In contrast, we demonstrate that maximum parsimony methods for lineage tracing data not only underestimate branch lengths, but also yield branch lengths which are not proportional to time, due to the nonlinear decay in the number of mutations on branches further from the root. On lineage tracing data from a mouse model of lung adenocarcinoma, we show that LAML infers phylogenetic distances that are more concordant with gene expression data compared to distances derived from maximum parsimony. The LAML tree topology is more plausible than existing published trees, with fewer total cell migrations between distant metastases and fewer reseeding events where cells migrate back to the primary tumor. Crucially, we identify three distinct time epochs of metastasis progression, which includes a burst of metastasis events to various anatomical sites during a single month.

**Software:** https://github.com/raphael-group/LAML

**Data availability:** https://github.com/raphael-group/laml-experiments

## 1 Introduction

Deriving the history of cell division and differentiation events that produce a multicellular organism from an embryo is of fundamental importance in developmental biology [26, 7, 28, 47, 44, 35] and tumor development [31, 6, 52, 43]. Since the cell division process is typically not directly observed, one may attempt to infer the history of cell divisions using heritable markers (e.g. somatic mutations) measured in cells at the present time. Such *retrospective lineage tracing* is challenged by the infrequency of naturally occurring somatic mutations which provide insufficient signal to construct a cell phylogeny. Recently, *dynamic lineage tracing* has overcome this challenge by using genome editing technologies, such as CRISPR/Cas9, to induce heritable mutations at pre-defined loci (i.e. *target sites* [29]) in the genomes of growing cells. These target sites are then sequenced via single-cell sequencing (sc-Seq) of a population of descendant cells to read out the induced mutations [28, 46]. Crucially, dynamic lineage tracing allows control over the strength of phylogenetic signal present in a cell culture by adjusting the number of target sites and genome editing parameters (e.g. amount of guide RNAs in CRISPR/Cas9). With suitable calibration, sufficient phylogenetic signal can be obtained for accurate reconstruction of cell lineage trees. While many dynamic lineage tracing technologies have been developed [28, 17, 35, 44, 2, 10, 5, 23, 52, 7, 9], we focus on more advanced technologies that are capable of recording medium to long experiment time, such as CARLIN [5], iTracer [23], Chan et. al [7], and Yang et. al [52].

Multiple computational methods have been developed to construct a cell lineage tree from dynamic lineage tracing data, including Cassiopeia [25], DCLEAR [21], AMbeRland [8], Startle [39] TiDeTree [40], and others. Several of the earlier methods were benchmarked in a DREAM challenge [20]. We classify existing methods into four categories: *distance-based methods* [48, 21], *maximum parsimony methods* [25, 39], *maximum likelihood methods*[16, 53], and *Bayesian methods*[40]. While it is well known in species phylogenetics that maximum likelihood and Bayesian methods often outperform maximum parsimony and distance-based methods [22, 19, 45], current methods in these categories for dynamic lineage tracing data [16, 53, 40] make assumptions that are only appropriate for older lineage tracing technologies [28, 10] but not for more recent CRISPR-based lineage tracing technologies [7, 52].

Three challenges hinder the application of existing probabilistic methods to newer dynamic lineage tracing data. First, the decay in the number of available target sites - a unique feature of the CRISPR-based editing system - is overlooked in many computational methods. In CRISPR-based lineage tracing systems, each target site accommodates at most one mutation, after which it can no longer be edited (i.e. *non-modifiable*). Consequently, the number of available (i.e. unmutated) target sites decreases over time, until mutations have saturated all available target sites, marking the end of *recordable time*. While innovations in experimental design have extended the recordable time of dynamic lineage tracing systems, on the computational front there has been no study of *how target site decay affects lineage tree reconstruction*. Second, lineage tracing data contains missing entries from two distinct mechanisms: *heritable missing data*, which results from silencing during the CRISPR-editing process and stochastic *dropout*, a technical error which occurs during single-cell sequencing. To the best of our knowledge, no existing computational method distinguishes these two types of missing data and leverages the phylogenetic signal from heritable missing entries. Third, current probabilistic methods do not analyze lineage tracing data with heterogeneous target sites (i.e. distinct per-site alphabet) as input.

We present the first probabilistic model and maximum likelihood method that can infer the cell phylogeny from data generated by recent dynamic lineage tracing technologies (such as [7] and [52]). Our proposed Probabilistic Mixed-type Missing (PMM) model, captures all key features of dynamic lineage tracing: non-modifiability of CRISPR, decay of the number of mutations over time, mixed-type missing data, and heterogeneous target sites,. Our maximum likelihood method, LAML (Lineage Analysis via Maximum Likelihood), jointly estimates tree topology, time-scaled branch lengths, and other parameters of the PMM model. In each iteration, LAML performs two steps: an EM algorithm for parameter estimation and tree rearrangements for topology search. Notably, the EM algorithm we derived under the PMM model is more efficient than that of existing phylogenetic models (such as the complex but widely used General Time Reversible (GTR) model). The complexity of the algorithm is presented in two cases: in the case of no heritable missing data, we present a closed form solution, and in the general case, we present an efficient block coordinate ascent approach.

We show that LAML outperforms existing methods on both topology and branch length inference on both simulated data and a mouse model of lung adenocarcinoma. In addition, we observe that maximum parsimony methods face a crucial problem in branch length estimation: maximum parsimony methods suffer from systematic bias where branches further from the root are more severely underestimated due to the decay in the number of unmutated target sites through time. Importantly, we demonstrate that LAML’s ability to estimate branch lengths in time units enables a new application for inferred cell phylogenies: we can track metastasis progression over time, from which we discover temporal epochs of metastasis progression in the lung adenocarcinoma model with distinct migration patterns.

## 2 The PMM model

Consider a dynamic lineage tracing experiment with *K* target sites^1^ and *N* sequenced cells. We refer to the data obtained from that experiment as the *observed character matrix*, denoted by **D**, which is an *N* ×*K* matrix. Entries in column *k* of **D** take values in 𝒜^(*k*)^ = {?, −1, 0, 1, …, *M* ^(*k*)^}, where 𝒜^(*k*)^ is the *alphabet* of target site *k*, 0 is the *unmutated state*, −1 is the *silent state*, ? is the *missing state*, and 1, …, *M* ^(*k*)^ are *mutated states*.

We represent the cell lineages by a rooted tree topology *T*. Each leaf node of *T* represents a sampled cell which corresponds to one row of **D** and the internal nodes represent unobserved ancestral cells. Let *ℒ*_*T*_ *𝒱*_*T*_, and *ℰ*_*T*_ be the *set of leaves, set of nodes*, and *set of edges* of *T*, respectively. Let *r*_*T*_ be the root of *T*. We assume that *r*_*T*_ has exactly one child (the progenitor cell needs time to divide) and all other internal nodes of *T* have exactly two children. Let (*u, v*) be the edge^2^ in *ℰ*_*T*_ from *u* to its child *v* (where *u, v* ∈ *𝒱*_*T*_). Each branch (*u, v*) of *T* has an associated branch length that shows the distance between *u* and *v*, which can be measured in either time units or mutation units, as will be described in section 2.1.

Given the character matrix **D**, our goal is to construct a cell lineage tree topology *T* and find all of its branch lengths that best describe **D**. We will solve this problem using a probabilistic approach: first, we describe a model for the likelihood 𝒫(**D**|*T*, **Θ**) where **Θ** includes the branch lengths and other parameters, and second, we develop an algorithm to find *T* that maximizes the likelihood. Our proposed Probabilistic Mixed-type Missing (PMM) model, consists of two layers corresponding to CRISPR/Cas9 editing of target sites and single-cell sequencing. Below we describe these two layers in more detail.

### 2.1 Layer 1: model of CRISPR/Cas9-induced mutations and heritable missing

Layer 1 describes the CRISPR/Cas9 editing process. At the beginning of the experiment, we assume there is exactly one *progenitor cell* that has all *K* target sites in the *unmutated state*. During the CRISPR/Cas9 process, each target site *k* either mutates into one of the *mutated states* in {1, …, *M* ^(*k*)^} or into the *silent state*. Recall that the cell phylogeny is denoted by *T*. We assume each site *k* mutates following a CTMC with the following transition rate matrix **Q**^(*k*)^:

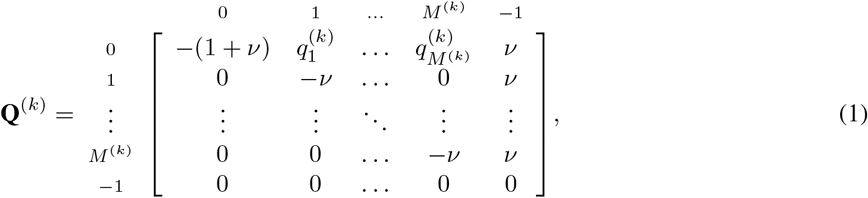

where 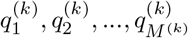 are the rates of transitioning from 0 to the mutated states of site *k* (i.e. 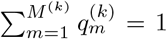 and *ν* is the heritable missing rate, which is shared across all sites.

Let *λ* be the editing rate (i.e. rate of changing from 0 to a mutated state) and *t*_*e*_ be the length of a branch *e* ∈ *ℰ*_*T*_ in time units. We assume *λ* is shared across all sites and branches and *t*_*e*_ for each branch *e* is shared across all sites. On an edge *e* ∈ *ℰ*_*T*_ at site *k*, the transition probabilities of every pair of states are determined by the *transition probability matrix*, 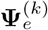, computed from **Q**^(*k*)^ as 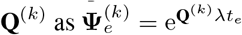. Performing matrix exponentiation gives the following:

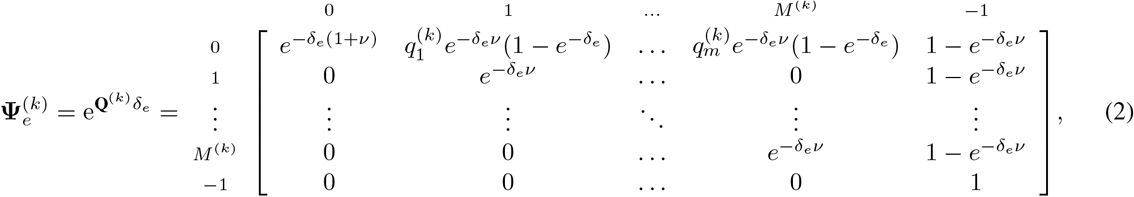

where *δ*_*e*_ = *λt*_*e*_, which is the length of *e* in mutation unit. Note that given *λ, t*_*e*_ and *δ*_*e*_ can be trivially computed from one another. In other words, let 𝕏 be the *random* matrix of size |*𝒱*_*T*_ | × *K* whose entries take values in 𝒜^(*k*)^ \ {?}.

Each entry in column *k* of 𝕏 represents the state of the corresponding cell at target site *k before single-cell sequencing*. Then for all *α*_*u*_, *α*_*v*_ ∈ 𝒜^(*k*)^ \ {?}, we have:

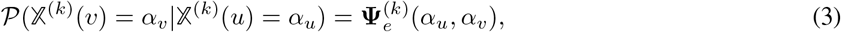

where 𝕏^(*k*)^(*u*) and 𝕏^(*k*)^(*v*) denote the entries of 𝕏 corresponding to site *k* and cells *u* and *v*, respectively.

The model is illustrated in Figure 1. Note that we assume that *t*_*e*_, *δ*_*e*_, and *ν* are shared for all sites, but allow each site *k* to have a distinct alphabet 𝒜^(*k*)^ and a distinct set 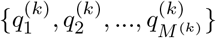. To avoid overparameterization, we treat all 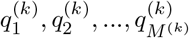 as hyperparameters (also referred to as *indel priors* in previous works [25, 39]). The user can specify these hyperparameters to be identical (as in [40]), or use external data to provide pre-estimated values (as in [25]). Notably, the transition matrix 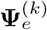 is sparse and has a special structure: non-zero entries are only in the first row, last column, and main diagonal.

**Figure 1.**
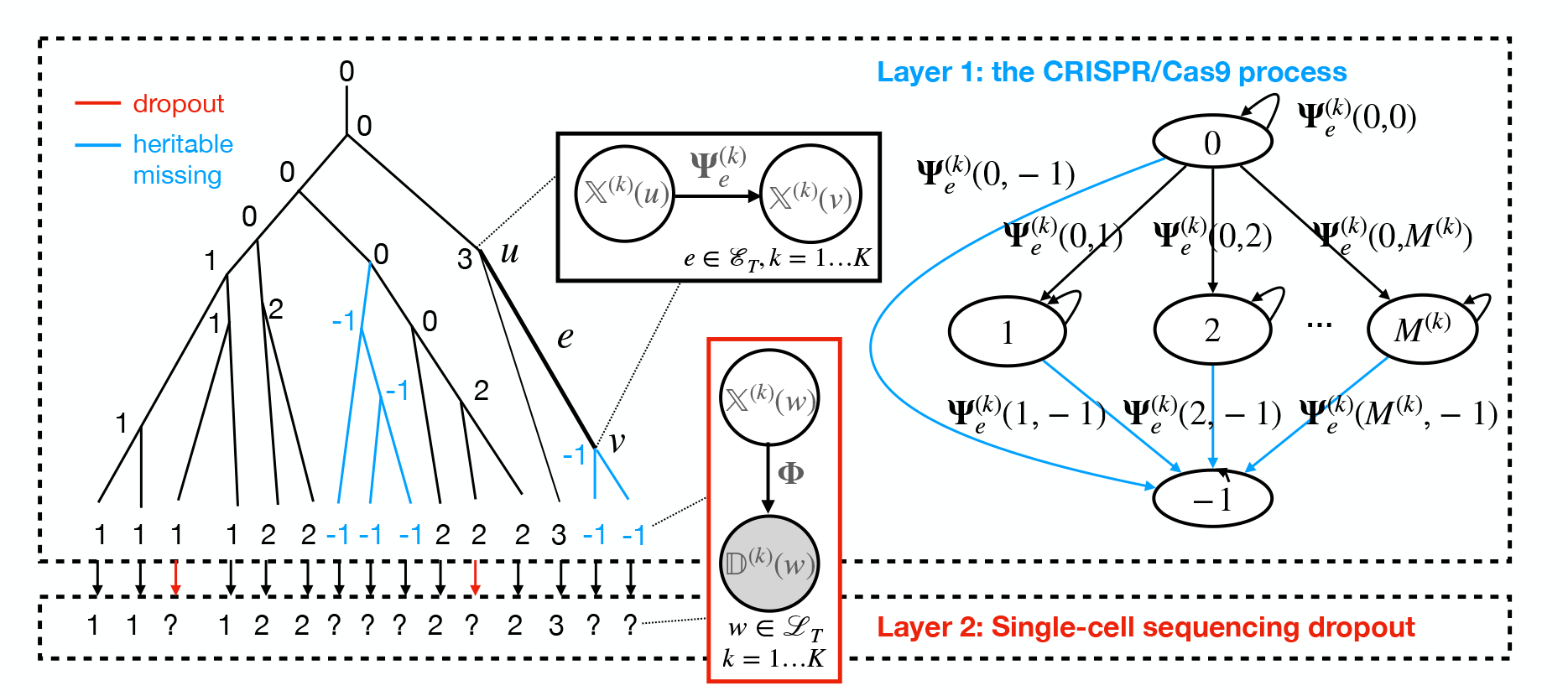
The PMM model with its two layers. Layer 1 (Top): A CTMC that models the process of CRISPR/Cas9 editing and heritable missing and (2) Stochastic dropout in single-cell sequencing. 0 denotes the unmutated state, −1 denotes the silent state, and other integers denote mutated states. State transitions are demonstrated by both a plate diagram and a transition graph. Transition probabilities are parameterized by **Ψ** (see Eq.2). Layer 2 (Bottom): A model for stochastic dropout in single-cell sequencing. The model is parameterized by **Φ** as indicated by the plate diagram (see Eq. 4). Observed missing data is represented by “?” and is a mixture of heritable missing (part of Layer 1) and dropout (part of Layer 2). The two types of missing data are indistinguishable in the observed data.

### 2.2 Layer 2: model of dropout from single-cell sequencing

The second layer of the model describes the stochastic generation of dropout during single cell sequencing. Recall that after single cell sequencing, characters in the silent state (i.e.−1) will be observed in the missing state with probability 1, while the other states have some probability to transition to the missing state depending on the dropout rate of single cell sequencing. We use *ϕ* to represent the *dropout rate* and represent the transition probabilities during single cell sequencing by the following matrix:

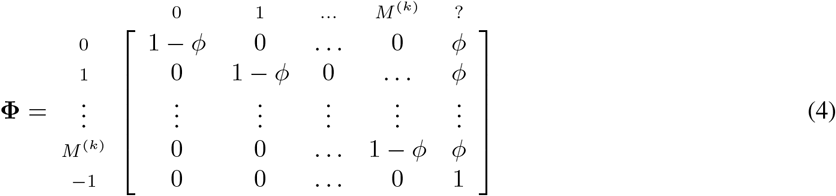

Let D be the *random* matrix of size |*ℒ*_*T*_| × *K* whose entries take values in *A* ^(*k*)^\ {−1}. Each entry in column *k* of 𝔻 represents the state of the corresponding cell at target site *k after single cell sequencing*. Note that **D**, the character matrix, is a realization (observed value) of D. For all *α* ∈ 𝒜^(*k*)^ \ {?}, *β* ∈ 𝒜^(*k*)^ \ {−1} and *w* ∈ *ℒ*_*T*_, we have:

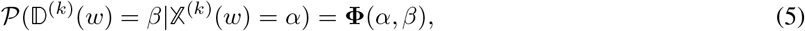

where 𝕏 ^(*k*)^(*w*) and 𝔻 ^(*k*)^(*w*) denote the entries of 𝕏 and 𝔻 corresponding to site *k* and cells *w*, respectively.

In summary, the PMM model has the following parameters: all branch lengths, the mutation rate *λ*, the silencing rate *ν*, and the dropout rate *ϕ*. In the rest of this paper, we use **Θ** = {( *δ*_*e*_, *λ, ν, ϕ*) } to denote these parameters. The model is illustrated in Figure 1.

## 3 LAML: Maximum likelihood inference

We assume that sites mutate independently, and thus given tree topology *T* and **Θ** describing branch lengths and missing data parameters, the log-likelihood log *L*(*T*, **Θ**; **D**) of the observed data **D** under the PMM model is the sum of the log-likelihoods of the individual sites:

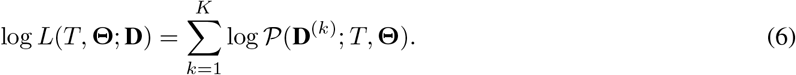

The probability 𝒫 (**D**^(*k*)^; *T*, **Θ**) of one site *k* is computed by marginalizing over all possible realizations of 𝕏 ^(*k*)^:

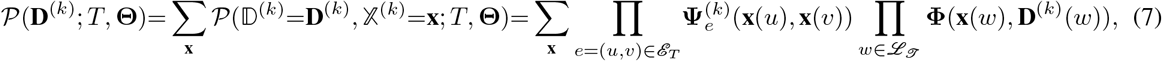

where **Ψ** and **Φ** are defined in Eqs. 2 and 4. While the sum in the above equation is over an exponential number of possible realizations of 𝕏 ^(*k*)^, the log-likelihood can be computed in linear time, using Felsenstein’s pruning algorithm [15]. See Supplementary Methods for a review of the pruning algorithm and the likelihood recurrences derived specifically for the PMM model.

### 3.1 Maximum likelihood inference

Our goal is to find the tree topology *T* and parameters **Θ** that maximize the log-likelihood function:

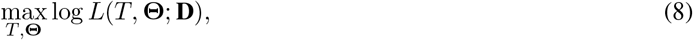

such that

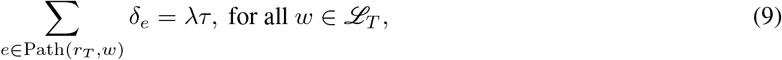

where Path(*r*_*T*_, *w*) denotes the path (a list of edges) from the root *r*_*T*_ to a leaf node *w*, and *τ* is length of the experiment (must be given by the user). Note that while **Θ** does not include {*t*_*e*_}, we can compute every *t*_*e*_ from *λ* and *δ*_*e*_ using *δ*_*e*_ = *λt*_*e*_. As such, we infer *branch lengths in both mutation units and time units*.

In general, maximum likelihood inference of phylogenetic trees is NP-hard [36]. Therefore, we use the following algorithm to find the maximum likelihood tree and **Θ**, iterating over two subroutines: (i) for a fixed tree topology *T*, find **Θ** to maximize *L*(**Θ**; *T*, **D**), and (ii) search for the optimal tree topology *T*. To solve (i), we devise *an efficient EM algorithm* and to solve (ii) we use simulated annealing with Nearest-Neighbor-Interchange (NNI) operations. Details of both steps are provided below.

### 3.2 Estimation of Θ on a fixed tree topology *T*

Given a tree topology *T* the maximum likelihood estimate 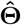 is defined as

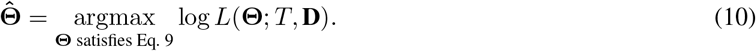

We compute 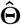 using the expectation maximization (EM) algorithm. Note that since *T* is fixed, we omit *T* from the equations below for brevity.

#### 3.2.1 The EM algorithm

The expectation maximization (EM) algorithm has been used to compute maximum likelihood estimates of branch lengths given a fixed tree topology [18, 42], which iteratively updates 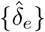 by alternating between an E-step and M-step. Building on these works, we derive an efficient EM algorithm to compute the maximum likelihood estimates for all parameters 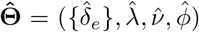, which are the estimated branch lengths 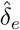, mutation rate 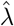, heritable missing rate 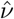, and dropout rate 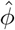. This algorithm leverages the special structure of the transition matrix (equation (4)) to reduce the complexity of the E-step and simplify the optimization problem involved in the M-step.

We briefly describe the E-step and the M-step below with the full proof and pseudocode in Supplementary Methods.

**E-step** At iteration *t* + 1, in the E-step we compute the expected values of the transitions 𝕏 (*u*) → 𝕏 (*v*) for every edge *e* = (*u, v*) of *T* and expected values of the transitions 𝕏 (*w*)→ 𝔻 (*w*) for every leaf *w* with respect to the estimate 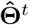 of **Θ** at iteration *t*. Because both **Ψ**_*e*_ and **Φ** are sparse and have special structure, we group the transition types of 𝕏 (*u*) → 𝕏 (*v*) into the following 5 groups: (i) 0 to 0 (*z* → *z*), (ii) 0 to any mutated state (*z* → *a*), (iii) 0 to the silent state -1 (*z* → *m*), (iv) any mutated state to another mutated state *a* → *a*, and (v) any mutated state to silent state -1 (*a* → *m*). Similarly, we group the transition types of 𝕏 (*w*) → 𝔻 (*w*) into the following two groups: no-dropout *ℬ* and Dropout 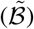. The expected values of these transition types are computed as follows:

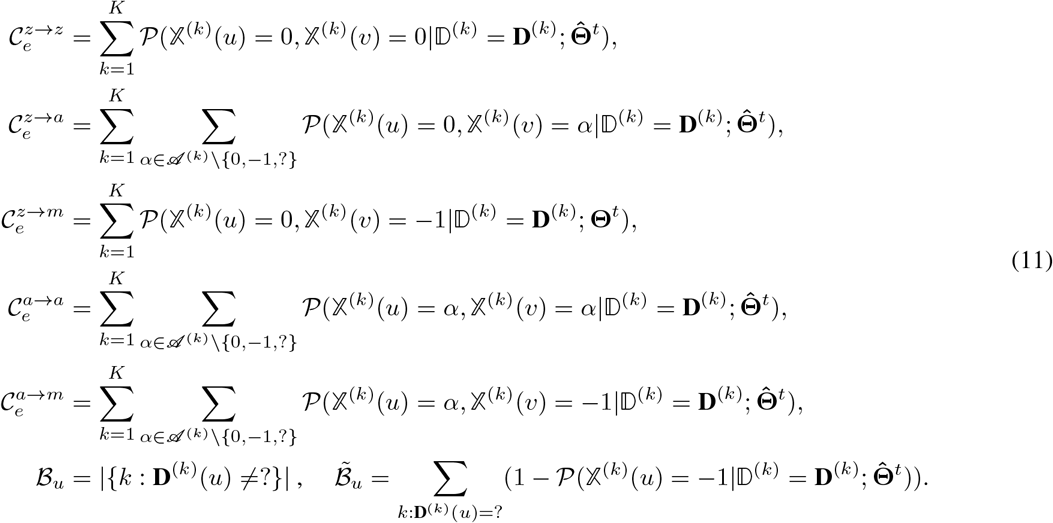

It is known that all the posterior probabilities 𝒫 (𝕏 ^(*k*)^(*u*) = *α*_*u*_, 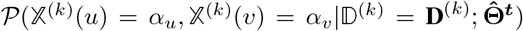) can be computed in 𝒪 (*MNK*) [18, 42] for all combinations of *e* = (*u, v*), *α*_*u*_, *α*_*v*_, where *M* is the maximum alphabet size, *N* is the number of sequenced cells, and *K* is the number of target sites. Therefore, the complexity of the E-step is also 𝒪 (*MNK*). While this complexity is acceptable for most current lineage tracing data, we further reduce the complexity to 𝒪 (*NK*) by leveraging special properties of our model to compute the expected values without knowing all the posterior probabilities (details in Supplementary methods).

**M-step** The M-step of iteration *t* + 1 is the following optimization problem:

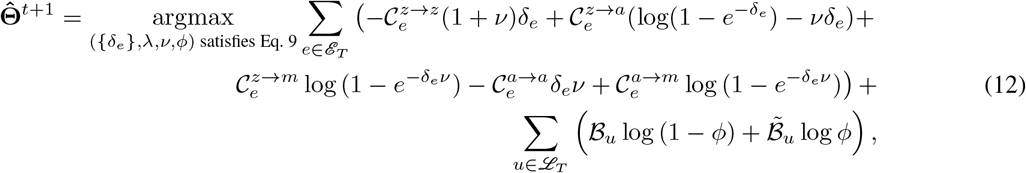

where the coefficients (of form 𝒞 and ℬ) in Eq. 12 depend on 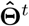 and were computed in the E-step. While this problem is not convex in general,in a special case where the heritable missing rate *ν* = 0, the problem is convex and if we ignore the constraints, we can derive a closed-form solution by a simple change of variables. When *ν* ≠ 0, the problem is still solved efficiently using block coordinate ascent.

##### Case 1 heritable missing rate

*ν* = 0. If *ν* = 0, then 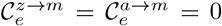 for all edges *e*. In addition, for all *u* ∈ ℒ_*T*_ we have 𝒫(𝕏^(*k*)^ = 0), so 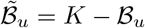. Thus, the M-step described by Eq. (12) reduces to:

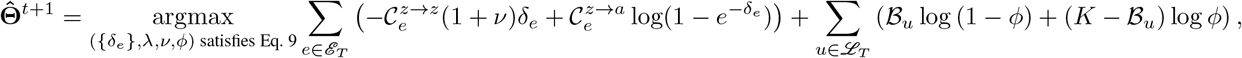

This optimization problem is convex (the objective function is separable, its second partial derivatives are negative, and the constraints are linear).

**The closed-form M-step when** *ν* = 0 **and ignoring constraints** *If we ignore the constraints given in Eq. 9*, we obtain a closed-form solution, as follows. Let 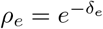. Then the problem reduces to

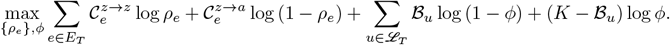

We solve the above problem by simply finding the roots of the partial derivatives, obtaining the following *closed-form* solution:

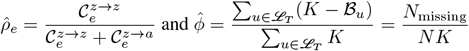

where *N*_missing_ is the total number of missing entries “?” in **D**. Then 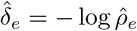. Because 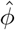 only depends on **D**, we compute 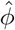 outside the EM algorithm. We note that because the constraints given in Eq. 9 are ignored, this closed-form formula only gives solution for branch lengths in mutation units and does not give a closed form solution for *λ*.

**Solving M-step in general case** If *ν* ≠ 0, the M-step problem is not always convex. However, the problem is convex separately with respect to *ϕ*, all *δ*_*e*_, and *ν*. In this case, we use block coordinate ascent to find a local optimum, where we sequentially optimize *ϕ, δ*, and *ν* (See Supplementary methods S1.6 for proof of convexity and the block coordinate ascent algorithm). In summary, our EM algorithm is an iterative algorithm where each iteration consists of two steps: (i) **E-step**: compute Eq. 11 and (ii) **M-Step**: solve for **Θ** in Eq. 12. We present Pseudocode in Algorithm box 1.

##### Algorithm 1

Pseudocode for the EM algorithm. Inputs: **D**: the character matrix, *T* : the tree topology, *ϵ*: the convergence threshold, and MaxIter (optional): the maximum number of EM iterations allowed (default to 1000). Output: the maximum likelihood estimate 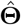.

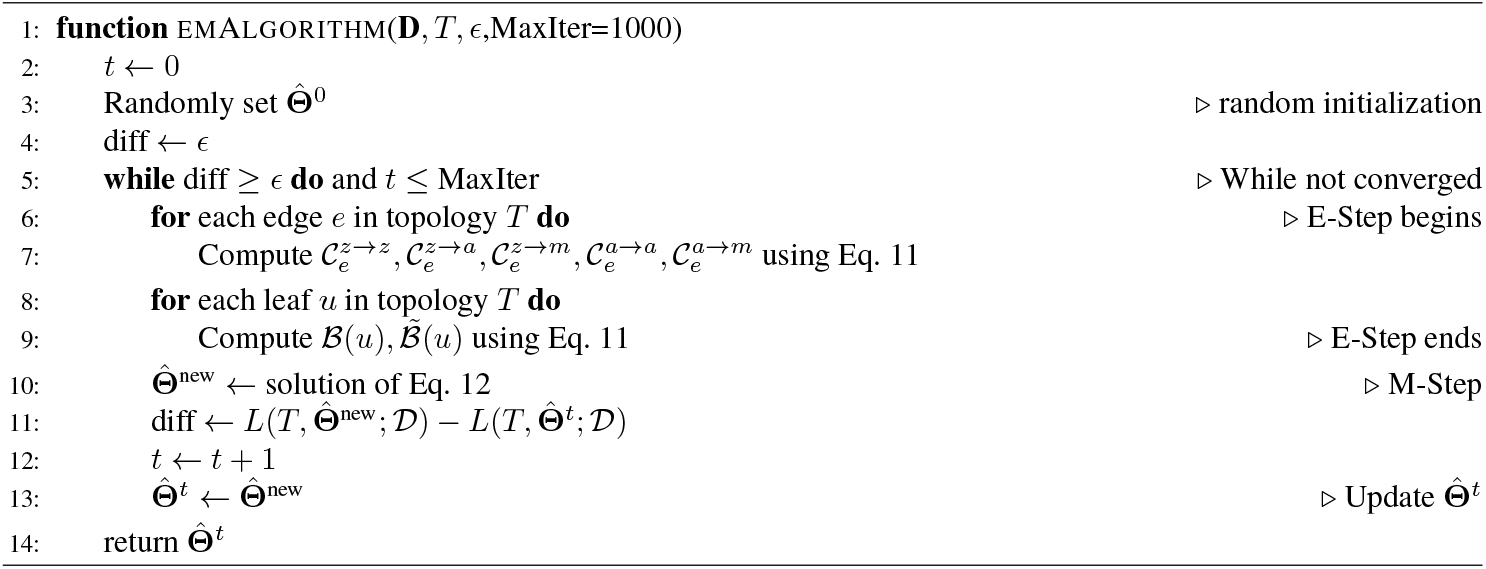

### 3.3 Topology search

We now describe Lineage Analysis via Maximum Likelihood (LAML), an algorithm that alternates between proposing a new tree topology and using the proposed EM algorithm to find the maximum likelihood estimate 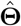 on this topology. Topology search is performed using nearest neighbor interchange (NNI) with simulated annealing. We provide the following pseudocode for LAML (Algorithm Box 2). See further details about simulated annealing in Supplementary methods S1.7.

#### Algorithm 2

Pseudocode for the LAML algorithm. Inputs: *D*: character matrix, *T*_0_: starting tree (either random or constructed using a fast tree construction method), *ϵ*_topology_ and *ϵ*_em_: the thresholds determining convergence of topology search and EM algorithm, respectively. Output: a tree topology *T* and parameters 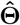.

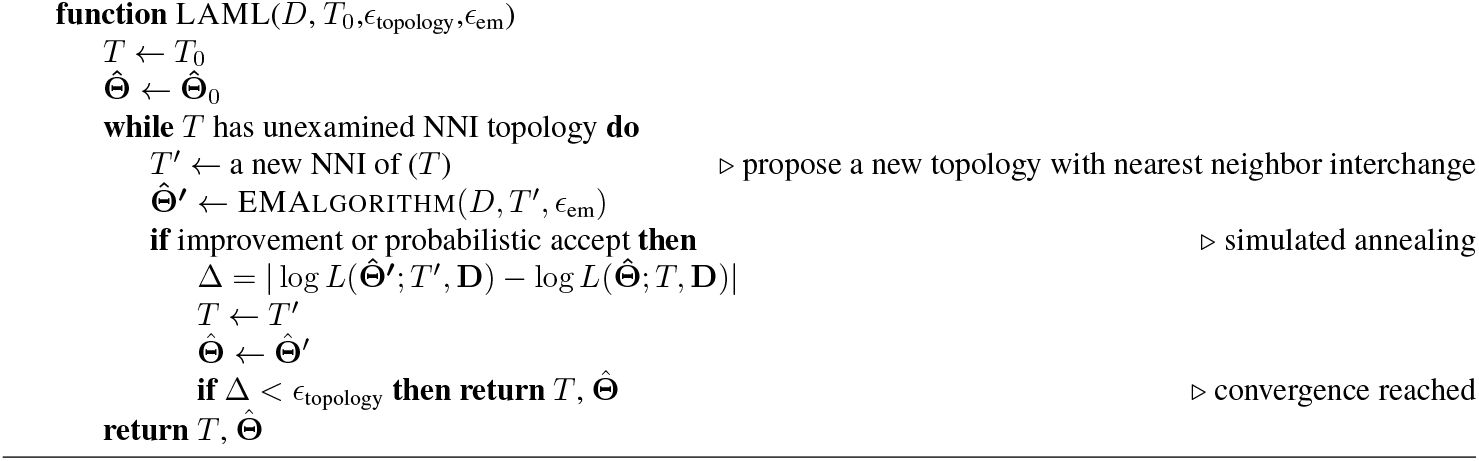

## 4 Results

We evaluate LAML on simulated data and two datasets from *in vivo* lineage tracing experiments: a mouse model of metastatic lung adenocarcinoma (KP-Tracer) [52], and a cellular developmental study of mouse embryos with true recorded cell lineages (intMEMOIR) [10].

### 4.1 Simulated data

We compare LAML to four other algorithms for constructing trees from lineage tracing data: the Cassiopeia-Greedy method [25], distance-based Neighbor Joining (implemented in Cassiopeia [25]), DCLEAR [21] – another distance-based method, and Startle-NNI [39] – a maximum-parsimony method. We evaluate these methods on simulated trees with 1024 leaves (i.e. 10 cell generations) generated using the birth-only process in the Cassiopeia simulation engine [25]. Branch lengths in time units are drawn according to a lognormal distribution with mean 1 and standard deviation 0.1. We simulate *K* = 30 target sites under the PMM model, with the values of hyperparameters *q*^(*k*)^ at each site *k* selected to match a recent lineage tracing dataset of trunk like structures (TLS) [4]. We also calibrate the simulated data to match two other statistics in the TLS dataset: the expected proportion (64%) of non-zero (mutated) entries in the character matrix and the expected proportion (25%) of missing data. To achieve 64% non-zero entries in the simulated data, we set the mutation rate *λ* = 0.095. To obtain a proportion of 25% overall missing data (with varying proportions of heritable and non-heritable missing data), we calibrate *ϕ* and *ν* for each of the five model conditions, as summarized in Table S2. To model the sampling of sequenced cells, each simulated character matrix is subsampled to 250 cells and the model tree is restricted to the same subset. We repeat the subsampling procedure five times for each character matrix. For each of five model conditions and ten sets of mutation alphabet hyperparameters, we created five replicates each, for a total of 250 replicates.

#### 4.1.1 LAML infers more accurate tree topology and branch lengths than other methods

We compare the cell lineage tree topologies output by the five methods using Robinson-Foulds (RF) error (i.e. normalized RF distance between true and estimated trees, defined in Supplementary data S2.1). Our results show that the trees estimated by LAML have the lowest RF error in all model conditions (Figure 2A). For all methods, RF error decreases as the proportion of heritable missing increases. For instance, LAML’s RF error decreases from 0.38 at h0d100 to 0.2 at h100d0. However, the improvement LAML achieves over other methods also increases with the proportion of heritable missing. For instance, compared to Startle – the second best method – the RF error of LAML is lower by 0.06 at h0d100 and lower by 0.14 at h100d0. This result suggests that LAML better leverages the phylogenetic signal in heritable missing data than the other methods.

**Figure 2.**
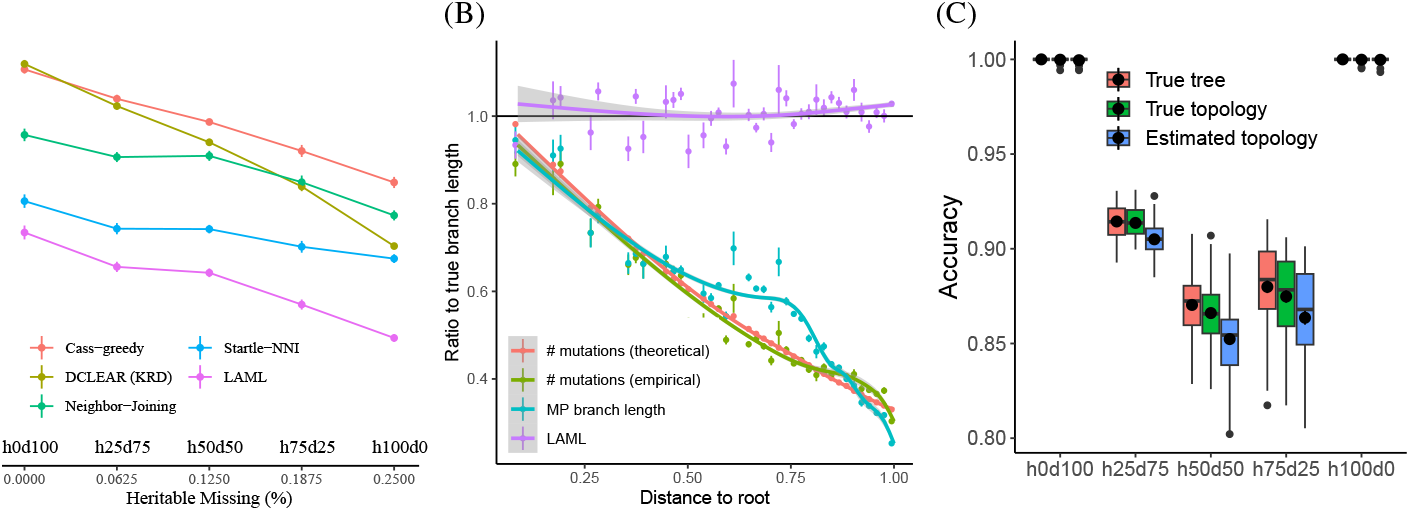
Comparison of LAML and other methods on simulated data. (A) Evaluation of tree topologies using normalized RF error under 5 different percentages of heritable (‘h’) and dropout (‘d’) of missing data. (B) The ratio of estimated to true branch lengths on true tree topologies Black line shows the perfect fit where estimate/true = 1. The *x*-axis is discretized into 50 bins and the estimated branch length by each method (or measure) is averaged for each bin, shown with error bar. (C) Classification accuracy for LAML in distinguishing the two types of missing data – heritable and dropout – shown for 3 scenarios: (red) True tree topology *T*_true_ with true parameters **Θ**_true_ given; (green) The true topology *T*_true_ is given, the MLE 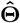 is estimated by LAML; (blue) both *T* and 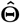 are estimated by LAML.

As a positive control, we also evaluate each method by its ability to optimize its own objective (Table S3 and Fig. S2). As expected, LAML has the highest log-likelihood, while Startle-NNI (a maximum parsimony method) typically has the lowest weighted parsimony cost. However, in 16% of the cases the tree inferred by LAML has a lower weighted parsimony cost than the Startle-NNI tree. One possible reason is because Startle-NNI is a heuristic algorithm that does not always find the true optimum and since there is a high correlation between weighted parsimony and log-likelihood, LAML will typically also produce trees achieving a good weighted parsimony score.

Next, we evaluate the accuracy of branch lengths estimated by LAML, where we give the method the true tree topology and use the EM algorithm to estimate branch lengths and missing data rates. Overall, the branch length estimates by LAML are accurate: the ratio of estimated to true branch length is around 1 regardless of that branch’s distance to the root (Fig. 2B, purple line), and there is a high correlation (*R* = 0.72) between the true branch lengths and LAML’s estimated branch lengths (Fig. S4). Furthermore, the estimated mutation rate 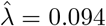 is very close to the true mutation rate *λ* = 0.095.

Although LAML is the only method that infers branch lengths in time units, we also made a best attempt comparison of LAML’s branch lengths to those inferred by other methods. Note that in order to perform this evaluation, we need a given method to be able to infer branch lengths for the *true tree topology*. Unfortunately, none of the 4 other software packages (Cassiopeia-Greedy, Cassiopeia-NJ, Startle, and DCLEAR) perform branch length inference on a given (i.e. fixed) tree topology. Therefore, we only benchmark against the *MP branch lengths* – standard practice to produce branch lengths on trees inferred by maximum parsimony methods, previously used for downstream analyses on lineage tracing data [52]. To compute MP branch lengths, we use Sankoff’s algorithm to infer the ancestral sequences and compute branch length as the number of mutations separating nodes from one another. We note that the MP branch lengths are not measured in time units; instead, each MP branch length is an estimate of the *number of mutations* that occurred on the corresponding branch (i.e. mutation count). As a best effort “conversion” of the MP branch lengths to time units, we divide them by the true mutation rate *λ* = 0.095 before evaluating them with respect to the true branch lengths. Overall, the MP branch lengths have much lower correlation to the true branch lengths (*R* = 0.39) than LAML branch lengths. More importantly, MP branch lengths also have a *systematic bias* where the ratios to the true branch lengths decrease with the distance from the root (i.e. branches further from the root are more seriously underestimated) (Fig. 2B, red and green lines).

We hypothesize that the observed systematic bias of MP branch lengths is the result of using mutation counts as a proxy to measure branch lengths. Because of the decay in the number of available target sites, *mutation counts are not proportional to branch lengths in time units* under the PMM model. Instead, the ratio of mutation counts to time-unit branch lengths actually degrades over time. To test this hypothesis, we compute both the theoretical and empirical mutation counts under the PMM model and added them as the two alternative means for branch length computation. Under the PMM model, the theoretical expected mutation count of each branch *e* = (*u, v*) is 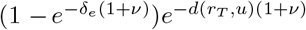 where *d*(*r*_*T*_, *u*) denotes the distance from the root *r*_*T*_ to *u* (we omit a proof here, but the derivation is straight-forward using Eq. 2). The empirical mutation counts are simply collected from simulated data. We also divide both the theoretical and empirical mutation counts by *λ* to “convert” them to time units. Consistent with our hypothesis, we observe that both the theoretical and empirical mutation counts are systematically biased if used as branch lengths, and that branches further from the root are more seriously underestimated (Fig. 2B, red and green lines).

#### 4.1.2 LAML accurately infers heritable missing and dropout rates

LAML produces accurate estimates for the missing data rates on all five model conditions, and furthermore these estimates are robust against topological error (Fig. S1A). However, more accurate tree topologies help reduce errors in these estimates. For instance, the values we estimate on the LAML tree topology (which achieved the lowest RF error) have the lowest RMSE. Note that LAML is the only method that estimates both heritable missing and dropout rates, so we cannot compare to other methods on this aspect.

We additionally evaluate LAML’s ability to distinguish heritable missing data and dropout. LAML achieves nearly perfect classification accuracy when all missing entries (which account for 25% of the whole data) arise solely from dropout (heritable: 0%, dropout: 100%, denoted as: h0d100) or solely from heritable missing data (heritable: 100%, dropout: 0%, denoted as: h100d0). However, the accuracy drops when there are mixed-type missing data, with the lowest accuracy (88% on true tree, 87% on true topology with estimated branch lengths, and 85% on estimated topology) observed in model condition h50d50 (Fig. 2C). Interestingly, the classification accuracy is higher with lower variance on data where heritable missing events account for fewer of the missing entries (h25d75), compared to when heritable missing events account for more of the missing entries (h75d25), suggesting that LAML correctly classifies more dropout missing events than heritable missing events, possibly due to an overestimation of dropout. In all model conditions, LAML’s missing data imputation is robust against errors in tree topology and branch length (≤ 2% difference when using the true versus estimated trees).

### 4.2 Mouse lung adenocarcinoma (KP-Tracer)

We analyze a dataset of lung adenocarcinoma metastasis in a mouse model [52]. We show below that LAML not only infers a cell lineage tree that is *more plausible* than the published trees inferred by Cassiopeia [52] and Startle [39], but also our time-resolved branch lengths enable the study of *metastasis progression*, a novel application of the cell lineage tree.

#### 4.2.1 Analysis of branch length estimation on six representative samples

We first use LAML to estimate branch lengths for the published tree topologies (inferred by Cassiopeia) for six representative samples (3432 NT T1, 3435 NT T4, 3520 NT T1, 3703 NT T1, 3703 NT T2, and 3703 NT T3) that have a high number of target sites (ranging from 29 to 73) and moderate numbers of cells (29 to 294). Note that in this experiment we fixed the tree topology and only estimate branch lengths. We compare the branch lengths estimated by LAML to the MP branch lengths (using Sankoff’s algorithm to estimate ancestral states and computing mutation counts on each branch, which are treated as branch lengths on maximum parsimony trees). We observe that LAML’s branch lengths are not proportional to the MP branch lengths (Fig. 3A), especially near the leaf nodes where many long LAML’s branches have a low mutation count by maximum parsimony (red branches). The ratio of the maximum parsimony branch length to the branch inferred by LAML (MP/LAML) decreases with the distance to the root (Fig. 3B). Such a pattern is consistent with the results shown on simulated data (Fig. 2B) where the MP branch lengths decrease with distance to the root while LAML branch length estimates do not, and is observed in all of the six tested samples (Fig. S8A in Supplementary). Inspired by earlier studies [52, 53], we use mutual information between gene expression distances and the phylogenetic distances obtained from estimated branch lengths as an evaluation metric for branch length estimation. We use this metric to benchmark the branch lengths inferred by LAML against the MP branch lengths and “topology distances” - a baseline method that assigns a unit length to every branch. We find that LAML’s phylogenetic distances yield substantially higher mutual information than the counterparts (Fig. 3C), and the same pattern is observed when testing on all six samples (Fig. S8B in Supplementary). Further details about data processing and evaluation metrics are in Supplementary data.

**Figure 3.**
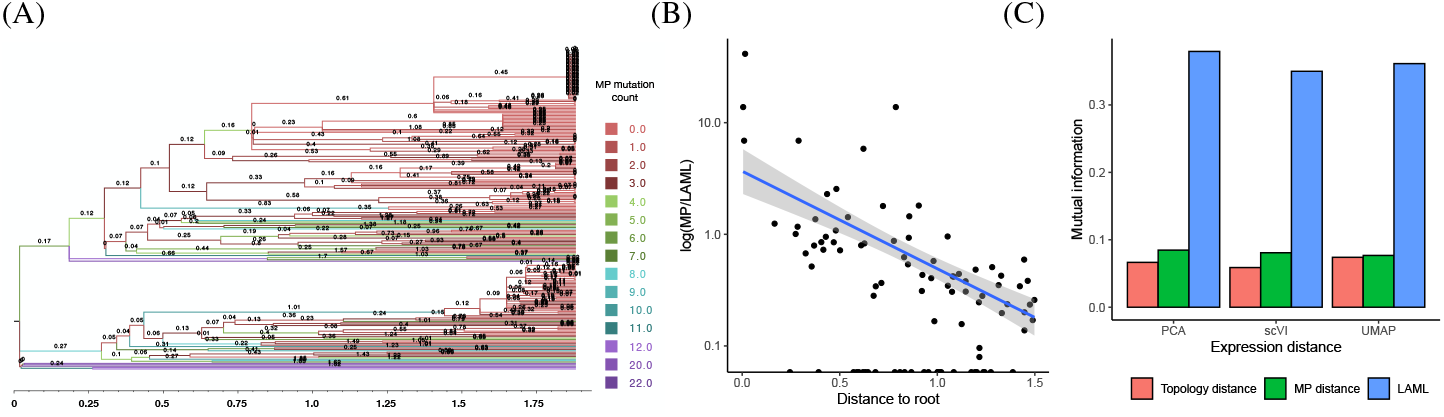
Evaluation of branch lengths on the cell lineage tree for sample 3432 NT T1 from the KP-Tracer mouse lung adeno-carcinoma lineage tracing study. (A) The tree topology published in [52] with branch lengths estimated by LAML. Numbers on branches are the lengths estimated by LAML and colors correspond to the number of mutations inferred by maximum parsimony. (B) The log-ratio between the length of a branch estimate by maximum parsimony and LAML versus the distance between the root node and the end of the branch. (C) Mutual information between gene expression and phylogenetic distances. Gene expression distance is computed in latent spaces computed by PCA, scVI, or UMAP. Phylogenetic distance computed by the sum of branch lengths between leaves using branches of length one (Topology distance), branch length estimates from LAML, or branch lengths from maximum parsimony (MP distance).

#### 4.2.2 Analysis of lineage tree topology on the largest sample

We use LAML to simultaneously infer the lineage tree topology and time-resolved branch lengths of the largest sample of KP-Tracer. This sample (3724 NT T1 All) has 21, 108 cells in total, but because many of the cells have identical CRISPR/Cas9 sequences, we only have 1,461 unique cells after deduplication. The tree inferred by LAML on the deduplicated data is shown in Fig. 4A. Interestingly, LAML produces a very different tree topology from the published trees inferred by Cassiopeia-Hybrid [52] (normalized RF 0.81) and Startle-NNI [39] (normalized RF 0.58). We discuss the biological implications of these different inferred topologies in this section and the next.

**Figure 4.**
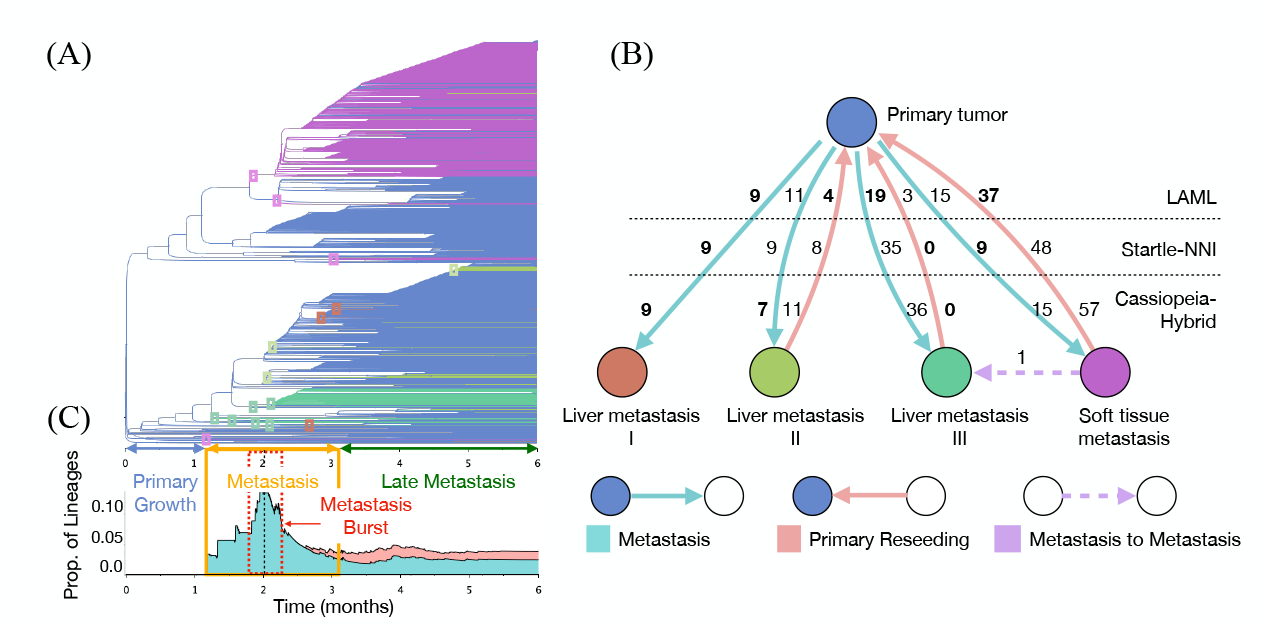
Migrations between anatomical sites in mouse lung adenocarcinoma sample 3724 NT T1 All. (A) Migration graph inferred from the de-duplicated trees estimated using Cassiopeia-Hybrid, Startle-NNI, and LAML. Nodes represent the anatomical sites, arrows represent migration events, and numbers represent number of migration events inferred from each estimated tree. Bold indicates the lowest number for each migration event. Note that the Metastasis to Metastasis event from Soft tissue to Liver III appears in all three methods. (B) LAML’s estimated phylogeny on 1,461 cells (de-duplicated data). Time-scaled branches are colored by the inferred anatomical location of incident child node. Small boxes indicate metastasis events. Time is divided into three epochs: Primary Growth (blue), Metastasis (yellow), and Late Metastasis (green). Red dashed lines mark the Metastasis Burst and the black dashed line marks the peak of metastasis burst. (C) Stacked area chart representing the proportion of lineages (branches) annotated as metastasis (aqua), reseeding (red) and metastasis to metastasis (purple) at each time point.

We benchmark the topology of LAML against Cassiopeia-Hybrid and Startle-NNI using the same procedure as in [39]: construct a cell *migration graph* [14] for each of these tree topologies and analyze these migration graphs. A migration graph is constructed as follows. Using the annotated anatomical location of the sampled cells and a tree topology, we first infer the anatomical locations of the ancestral nodes using Sankoff’s algorithm (as described in [14, 39]). Next, we collapse all branches that have parent and child nodes labeled identically. Finally, we merge all branches that have identical labels on both ends and use the multiplicity as edge weight. Thus, the migration graph constructed on a tree topology is a directed graph whose *nodes are anatomical sites, edges indicate migration events*, and *edge weights show the number of times a migration event happened* on that tree (Fig. 4A). Migration events are classified into three types: primary tumor to a non-primary anatomical site (*metastasis*), non-primary anatomical site to the primary tumor (*reseeding*), and non-primary anatomical site to a different non-primary anatomical site (*metastasis to metastasis*). Similar to [39], we define the *migration cost* as the sum of edge weights in the migration graph and use it as an evaluation metric for lineage tree topology. We also define the *reseeding cost* as the total number of reseeding events and use it as a secondary evaluation metric. See Supplementary Section S2.3 for further information.

Compared to the two published trees, LAML achieves both lower migration cost and lower reseeding cost (Table S4). In addition, the migration graph of LAML is more plausible than the two published trees (Fig. 4A), as it shows a clearly lower *reseeding to seeding ratio* (4 to 11) than Startle-NNI (8 to 9) and Cassiopeia-Hybrid (11 to 7) for liver metastasis II and a clearly lower total number of migration events for liver metastasis III (22 versus 35 versus 36). Interestingly, all tree topologies agree on a high number of reseeding events for soft tissue metastasis. This result is not surprising, however, as these events could have resulted from local spread of tumors prior to metastasis through the blood or lymphatic system to distant anatomical sites [41]. Nevertheless, the LAML topology is still more plausible as it yields the lowest number of reseeding events (37), followed by Startle-NNI (48), and finally Cassiopeia-Hybrid (57). All methods detect a single migration event from soft tissue metastasis to liver metastasis III, and no other migration events between other pairs of non-primary sites.

#### 4.2.3 Timing metastasis progression with time-resolved branch lengths

We use the time-scaled branch estimated by LAML to infer the time of cellular migrations spread between the primary tumor and the multiple anatomical sites. This analysis of the timing of metastasis was not performed in the original KP-tracer publication [52] because the lineage trees inferred by Cassiopeia do not have time-scaled branch lengths. Our results show three epochs of metastasis progression with distinct patterns of cell migration (Fig. 4B): Primary Growth (month 0-1), Metastasis (month 1-3), and Late Metastasis (month 3-6). In the Primary Growth epoch, cells grow and divide within the Primary tumor but do not migrate (i.e. no metastasis) (Fig. 4C). The Metastasis epoch starts at around month 1, with a gradual increase in the number of metastasis events and lasts until around month 2, followed by a sudden surge (i.e. Metastasis Burst) and a later gradual reduction until month 3 (Fig. 4C, aqua area plot). The first metastasis event is a soft tissue metastasis (around the end of month 1), consistent with previous reports of local spread of tumors in soft tissue [41] (Fig. 4C, red area plot). Notably, in month 2, there is a surge of metastases (i.e. the Metastasis Burst window, see Fig. 4C) that lasts about half a month (from month 1.75 to 2.25). We observe that during Metastasis Burst, the rate of metastasis (i.e. the number of metastases per cell lineage per month) increases from a background rate of 0.00349 to a rate of 0.11328, a greater than 30-fold increase, and during this short time there are metastases to multiple anatomical sites. After the metastasis burst, there is a gradual decrease in the number of migrations, until the beginning of month 3 when the number reaches a stable state. We mark this time point as the start of the Late Metastasis epoch. In Late Metastasis, there is a stable proportion of lineages with migrations at every time point. In addition, we also observe reseeding events during this epoch, where cells are inferred as migrating back to the primary tumor from distant metastases [14]. Interestingly, the ratio of reseeding to all migration events is maintained at a constant value of about 0.4 throughout the Late Metastasis epoch. Because there are a low number of unmodified target sites from month 3 forward (≤3), the phylogenetic signal is weaker during this epoch, and thus we expect higher uncertainty in the inference of both tree branch lengths and metastasis events.

Finally, we analyze the missing data rates inferred by LAML on this largest sample of the dataset. According to our estimates, this sample has mixed-type missing data: dropout rate *ϕ* = 0.043 and heritable rate *ν* = 0.010. Among the 753 missing entries in the data, LAML imputes 578 entries to be dropout and 175 entries to be heritable missing. The 175 heritable missing entries are attributed to 80 heritable missing events, with each event introducing 2 to 4 missing entries to the character matrix. Interestingly, the signal from heritable missing is lost when we estimate missing data rates and branch lengths using the Startle-NNI topology (*ϕ* = 0.057, *ν* = 0).

### 4.3 Mouse embryo cellular development (intMEMOIR)

We also apply LAML to a development study in mouse embryo which uses the intMEMOIR dynamic lineage tracing technology [10, 20]. The dataset has ground truth lineage trees collected through microscopic observation of the cell division process. The Bayesian MCMC algorithm TiDeTree, which also uses a continuous-time Markov model and allows heritable missing data, can be used to infer trees from data obtained with this dynamic lineage tracing technology. We compare LAML, TiDeTree, Startle-NNI and Cassiopeia on this dataset. LAML achieves the lowest average RF error compared to the ground truth tree of 0.53, followed by TiDeTree of 0.56, Startle-NNI of 0.58, and Cassiopeia of 0.63 (Figure S10A and Supplementary data S2.4). Interestingly, although the tree topologies inferred by the different methods have similar normalized RF distance from the true tree, they have large average RF distance from the LAML tree (Startle-NNI: 0.31, Cassiopeia: 0.46, TiDeTree (Pub): 0.46) (see Figure S10B). As the data has only ten heritable target sites and seems to support several very different tree topologies, there seems to be low overall phylogenetic signal, making lineage tree reconstruction difficult. Note that we were unable to run TiDeTree on either the simulated data or the KP-Tracer data above – both because the software does not scale to large numbers of cells and does not handle distinct per-site alphabets.

### 4.4 Runtime Evaluation

First, we explore the runtime of the LAML EM algorithm for branch length estimation. As supported by the theoretical complexity, in the special case where the heritable missing rate *ν* = 0, both the E-step and the M-step run in linear time (Fig. S5 black and yellow lines). Although we could not analyze the theoretical complexity of the M-step block coordinate ascent algorithm when *ν* ≠ 0, our empirical runtime is also close to linear (Fig. S5 grey line). However, the overall runtime of the EM algorithm, depends on the number of EM iterations needed (i.e. the convergence rate of the EM algorithm) and seems to be worse than linear (Fig. S6).

Next, we compare the overall runtime (the combined runtime of topology search and branch length estimation) of LAML to other methods. On simulated data (250 cells), Cassiopeia-Greedy and Neighbor Joining are the two fastest methods (Fig. S7). The runtime of LAML is comparable to Startle-NNI. Since we use the Startle-NNI topology to initialize LAML, we added the Startle-NNI runtime to the reported runtime of LAML. The average runtime difference between LAML and Startle-NNI is ≈ 100 minutes. We also benchmarked the runtime of LAML against TiDeTree on the intMEMOIR data (Fig. S11), which has a much smaller number of cells (≤77) and target-sites (10). Due to the long runtime of TiDeTree, we only ran it for 10 of the largest samples that have from 51 to 77 cells. Overall, we observe that LAML is much faster than TiDeTree: LAML finished in just under a minute while TiDeTree took ≈ 9 hours to infer the lineage tree of each sample.

## 5 Discussion

Single-cell lineage tracing technologies continue to advance in scale and resolution, increasing the need for accurate methods to infer lineage trees from the resulting data. We introduce LAML, a maximum likelihood method that jointly estimates tree topology, time-scaled branch lengths, dropout rate, and heritable missing rate under the newly developed PMM model for dynamic lineage tracing. LAML can be run to (i) estimate time-scaled branch lengths and missing data rates on a given tree topology, or (ii) estimate the tree topology as well as all parameters. We show on simulated data that LAML is more accurate for topology estimation than many existing methods, including a greedy approach (Cassiopeia-greedy), parsimony approaches (Cassiopeia-ILP, and Startle-NNI), and Neighbor-Joining (with multiple distance matrices, including DCLEAR). Importantly, while the simulation used parameters derived from real lineage tracing datasets, the data was generated under our proposed generative model (PMM), and so it not surprising that LAML - a maximum likelihood inference under this model - performs well. Additional simulations with model misspecification, correlation of heritable missing between consecutive sites, additional sc-Seq errors, variation of mutation rates among sites, or changes in mutation rate across time would be helpful. Future work should explore the robustness of LAML on these violations of the model and its performance in comparison to other methods.

We show that LAML produces more plausible lineage trees on a mouse model of lung adenocarcinoma [52]. We demonstrate that branch lengths inferred under a maximum parsimony model are systematically biased, as branches further from the root are more severely underestimated. Using the PMM model that realistically reflects the decay of mutation counts through time, we enable the estimation of branch lengths in time unit, which are needed in studying the dynamic of cell growth and migration through time. On the largest sample of mouse lung adenocarcinoma [52], we demonstrate that LAML can be used to time metastasis events, a novel application of the cell lineage tree. Importantly, LAML discovers an interesting pattern of metastasis progression which consists of 3 time epochs. The Primary Growth epoch spans about 1 month during which there is no metastasis event happens. The Metastasis epoch spans from month 1 to 3, during which time there is a surge of metastasis events. The Late Metastasis epoch starts at around month 3 when there is a stable proportion of lineages undergoing metastasis and Primary reseeding at every time point. There are multiple directions for future work. First, our biological findings will be strengthened by additional phylogenetic analyses such as bootstrapping [13] and relaxing the molecular clock assumption [24]. Second, the PMM model can be extended to model other aspects of cellular development, such as the presence of multiple progenitors, clustering of cell types and gene expression, and the spatial structure of the tissue or tumor. Additionally, the current PMM model does not incorporate the empirical observation that dropouts occur in blocks (that is, that consecutive target sites are dropped). Future work could aim to relax the assumption of independence across sites. Third, the sc-MAIL algorithm could be improved by further incorporating subtree-pruning-and-regrafting (SPR) moves through tree topology space. Finally, although LAML’s current runtime for estimation of the tree topology is acceptable scaling to 1, 461 cells on the largest sample from the KP-Tracer dataset, larger lineage-tracing datasets will likely be generated in the future. Both the runtime performance and scalability of LAML can be improved by parallelization [32], by phylogenetic placement, where one adds new sequences into an existing phylogenetic tree [50, 11], and algorithmically by a divide-and-conquer [49, 30] or hybrid framework, where one combines the accuracy of maximum likelihood with the speed of distance-based methods.

Our results show important implications for technological design and applications of dynamic lineage tracing. First, downstream analyses such as developmental tree inference, subclonal dynamics, and trajectory inference, benefit from the time-scaled branch lengths estimated by LAML. Second, experimentalists should aim for control over both the overall amount of missing data and the ratio of dropout to heritable missing types when developing new lineage tracing technology. Third, our probabilistic approach to tree inference could aid in the time-scaled reconstruction of *unobserved intermediate cell states* [33, 52]. Finally, more refined time calibration information, such as the fluorescence tagging in the lineage tracing vector [52, 44] or time points of multiple rounds of editing (present in technologies such as CARLIN, iTracer, scGESTALT), can be leveraged to better guide the conversion of the ML branch lengths to a time scale.

## Acknowledgements

This research was supported by NIH/NCI grant U24CA248453 to B.J.R. U.M. was funded by the Presidential Postdoctoral Fellowship at Princeton University. G.C. was funded by the NSF GRFP DGE grant 2039656. We thank Palash Sashittal and Henri Schmidt for help with running the Startle code; Sophie Seidel for help with running the TiDeTree code; and Michelle Chan for helpful discussion.

## S1 Supplementary methods

### S1.1 Extended notations

Consider a dynamic lineage tracing procedure consisting of *K* target-sites. At the beginning of the experiment, there is exactly one *progenitor cell* that has all target-sites in the *unmutated state*, denoted by 0. During the CRISPR/Cas9 process, each target-site *k* of any cell can either mutate into one of the *M* ^(*k*)^ *mutated states* in {1, …, *M* ^(*k*)^} or go to the *silent state*, denoted by −1. During the single-cell sequencing (sc-Seq) process, cells in the silent state cannot be determined and will show up in the *missing state*; in addition, cells in the other states can also turn into the missing state because of dropout. We denote the missing state by “?” and refer to the set 𝒜^(*k*)^ {= ?,−1, 0, 1, …, *M* ^(*k*)^} as the *alphabet* of the target-site *k*. We assume that there are *N* cells sequenced by sc-Seq, so the dynamic lineage tracing data consists of *N* × *K* entries. Following the convention in phylogenetics, hereafter, we refer to *target-site* as *site* for brevity.

We represent the cell lineage of the lineage tracing data by a rooted tree *T*. Let *ℒ* _*T*_, *𝒱* _*T*_, and *ℰ* _*T*_ be the *set of leaves, set of nodes*, and *set of edges* of *T*, respectively. Let (*u, v*) be the edge in *ℰ*_*T*_ from *u* to its child *v* (where *u, v* ∈ *𝒱*_*T*_), and *r*_*T*_ be the root of *T*. Unless otherwise specified, *e* is used to denote the edge connecting *u* to *v*. Throughout this paper, we use *edge* and *branch* interchangeably. We assume that the root of *T* has exactly one child and all of its other internal nodes has exactly two children. For each node *v* ∈ *𝒱*_*T*_, we let *T*_*v*_ be the clade of *T* rooted at *v, 𝒞*_*T*_ (*v*) be the leafset of *T*_*v*_, and 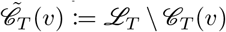. When the context is clear, the subscript *T* can be omitted for brevity.

Throughout this paper, we will use the following two conventions for random/realization matrices/vectors: (1) blackboard-bold letters (e.g. 𝔸, 𝕒, 𝔹, 𝕓, ℂ, 𝕔) will be used to indicate a *random* entity while normal bold letters (e.g. **A**,**a**,**B**,**b**,**C**,**c**) will be used to indicate a realization. (2) the uppercase letters (e.g. 𝔸, 𝔹, ℂ and **A**,**B**,**C**) are reserved for matrices while lowercase letters (e.g. 𝕒, 𝕓, 𝕔 and **a**,**b**,**c**) are reserved for vectors. For a set *ℐ* of *N* cells with *K* sites, we define the CRISPR/Cas9 sequences of *ℐ* by a *N* ×*K* matrix and refer to it as the *cell-state representation matrix* of *ℐ*, which has rows indexed by *ℐ* and columns indexed by the sites. We let 𝕏 be the cell-state representation random matrix of *𝒱*_*T*_ *before sc-Seq*, and **X** be its realization. Similarly, we let 𝔻 be the cell-state representation random matrix of *L*_*T*_ *after sc-Seq*, and **D** be its realization. Note that entries of **X** take values in 𝒜^(*k*)^ \{?} while entries of **D** take values in 𝒜^(*k*)^ \ {−1}. In dynamic lineage tracing, 𝔻 is observed exactly once and 𝕏 is hidden. Therefore, throughout this paper, **D** represents the lineage tracing data, which is also the sole *observed* realization of 𝔻. Borrowing the terminology from previous work [39, 52], we also refer to **D** as the *character matrix*.

We use the following conventions for any cell-state representation matrices, both random and realization. First, the superscript •^(*k*)^ is used to denote any entity associated with site *k*. For example, 𝕏 ^(*k*)^ denotes the *k*^*th*^ column of 𝕏. Second, we use •^(*k*)^(*v*) to refer to the element of the cell-state representation matrix associated with the row index *v* and column index *k*. For example, 𝕏 ^(*k*)^(*v*) denotes the random variable in 𝕏 that is associated with node *v* and site *k*. Finally, if **M** is a cell-state representation matrix and *ℐ* is a subset of the row indices of **M**, then we let **M**|_*ℐ*_ be the matrix constructed from the rows of **M** whose indices belong to *ℐ*. The same convention is applied to 𝕏, 𝔻, and their realizations. However, for the sole realization **D** of 𝔻, we let **D**_*v*_ be the shorthand for 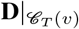 and 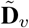 be the shorthand for 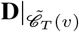.

**Table S1:**
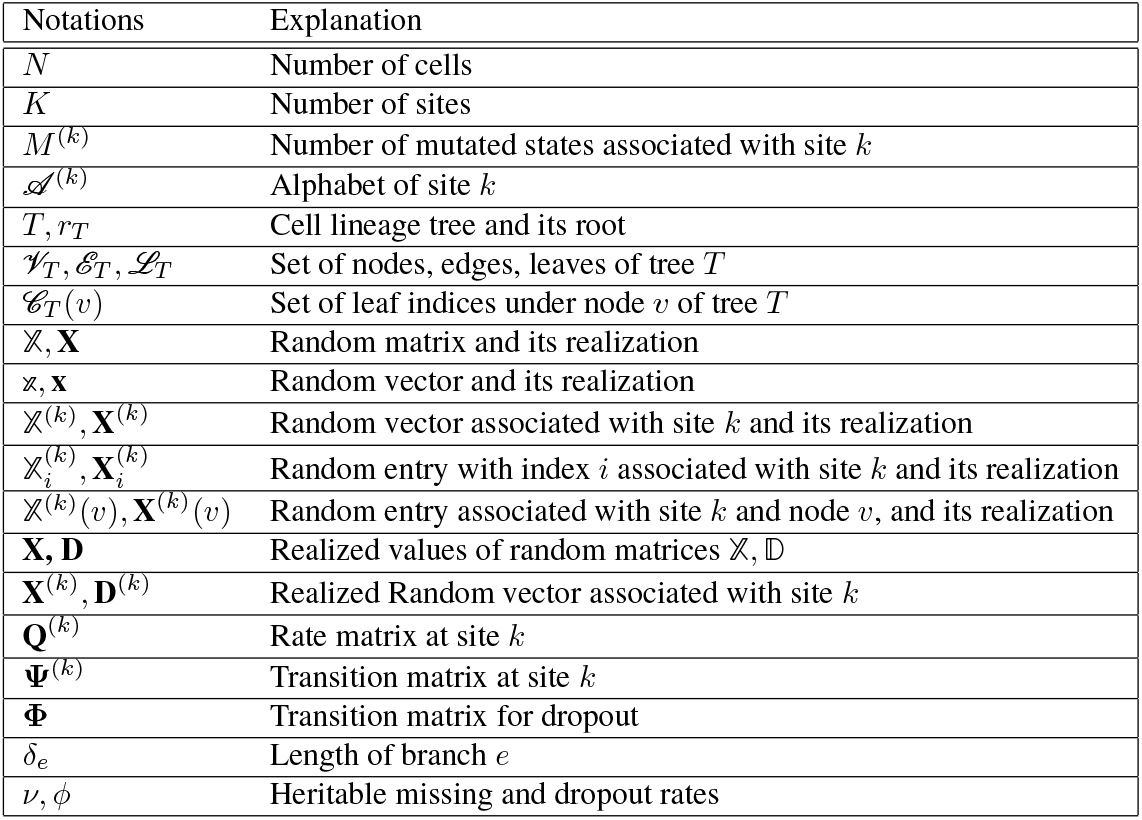
List of Notations.

### S1.2 Computing the log-likelihood given *T* and Θ

The log-likelihood can be computed in linear-time using Felsenstein’s pruning algorithm [15]. In this section we show that the pruning algorithm can be applied to our model by presenting recurrence equations.

To describe his algorithm, Felsenstein introduced the concept of *partial likelihood* at a node in the tree. Here we use the term **inside likelihood** (see Definition 1) to refer to the partial likelihood described by Felsenstein, for the purpose of distinguishing it from the **outside likelihood** that will be defined later. Consider a node *u* ∈*𝒱*_*T*_. The *node-inside likelihood of u* is defined as follows:

#### Definition 1.

*The node-inside likelihood (a*.*k*.*a partial likelihood) of u at site k w*.*r*.*t a realization α*_*u*_ *of𝕏* ^(*k*)^(*u*), *denoted by* 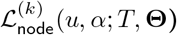, *is the likelihood of observing* 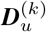 *given* 𝕏 ^(*k*)^(*u*) = *α*_*u*_. *In other words:*

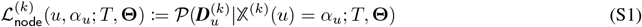

The node-inside likelihoods of all nodes *u* at all states *α*_*u*_ ∈𝒜^(*k*)^ can be computed in one bottom-up tree traversal, using the following recurrence:

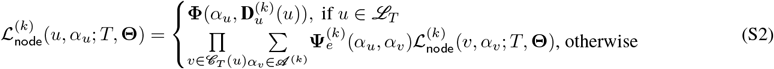

In the PMM model, 𝕏 ^(*k*)^(*u*) can only take state 0. Therefore, the full likelihood 𝒫 (**D**^(*k*)^; *T*, **Θ**) is simply the node-inside likelihood of the root node w.r.t. state 0. In other words, for all site *k* we have:

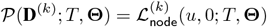

Thus, the full likelihood can be computed in one bottom up traversal. The complexity is therefore 𝒪 (*NKM*) where *N* is the number of cells, *K* is the number of target sites, and *M* is the maximum alphabet size.

### S1.3 The edge-inside likelihood, the outside likelihood, and the posterior

In this section, we define the **outside likelihood** and the **posterior** probabilities, which can be used later. We will show equations relating the inside likelihood, outside likelihood, and the posterior, and present a linear-time algorithm to compute all these entities for a tree (i.e. an extended version of the Felsenstein’s prunning algorithm). We note that similar equations have been previously described in the literature (see [42] for example). Below we show the definitions and recurrence equations, but skip the proofs for brevity.

#### S1.3.1 The edge-inside likelihood

We first define the *inside likelihood of a branch*, as follows:

##### Definition 2.

*The edge-inside likelihood of e* = (*u, v*) *at site k w*.*r*.*t. a realization α*_*u*_ *of* 𝕏 ^(*k*)^(*u*), *denoted by* 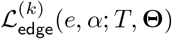, *is the probability of observing* 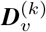 *given* 𝕏 ^(*k*)^(*u*) = *α*_*u*_. *In other words:*

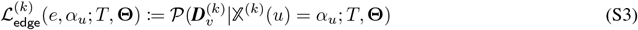

The edge-inside likelihood of *e* = (*u, v*) at any state *α*_*u*_ can be computed by summing over the inside likelihoods of *v* at all states *α*_*v*_, as follows:

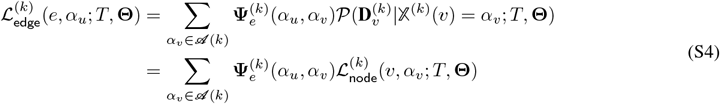

Using Eq. S4, we can extend the bottom up traversal to also compute the inside likelihoods at all branches.

#### S1.3.2 The outside likelihood

##### Definition 3.

*The outside likelihood of a node v at site k w*.*r*.*t. a realization α*_*v*_ *of* 𝕏 ^(*k*)^(*v*), *denoted by* 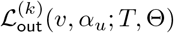, *is the joint probability of* 𝕏 ^(*k*)^(*v*) = *α and* 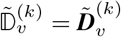. *In other words:*

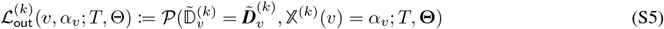

The outside likelihood can be computed using the following topdown recurrence:

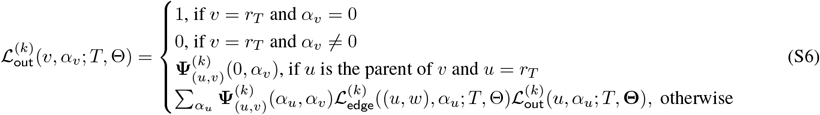

where in the last case, *u* and *w* are the parent and sister of *v*, respectively. In other words, one can compute the outside likelihood of any node from the outside likelihood of its parent and the inside likelihoods of the sister branch.

#### S1.3.3 The posterior

Next, we define the *posterior probability of a node* and *posterior probability of an edge*.

##### Definition 4.

*The posterior probability of a node u at site k w*.*r*.*t. the realization α*_*u*_ *of* 𝕏^(*k*)^(*u*), *denoted by* 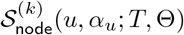, *is the probability of* 𝕏 ^(*k*)^(*u*) = *α*_*u*_ *given𝔻* ^(*k*)^ = ***D***^(*k*)^. *In other words:*

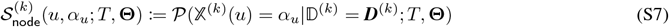

The posterior probability of a node can be computed using the inside and outside likelihoods, as follows:

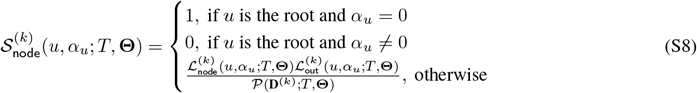

##### Definition 5.

*The posterior probability of an edge e* = (*u, v*) *at site k w*.*r*.*t. the realization α*_*u*_ *of* 𝕏 ^(*k*)^(*u*) *and α*_*v*_ *of* 𝕏 ^(*k*)^(*v*), *denoted by* 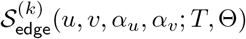, *is the joint probability of* 𝕏 ^(*k*)^(*u*) = *α*_*u*_ *and* 𝕏 ^(*k*)^(*v*) = *α*_*v*_ *given* 𝔻^(*k*)^ = ***D***^(*k*)^. *In other words:*

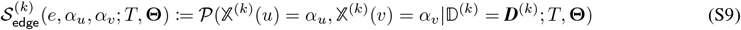

The posterior probability of an edge *e* = (*u, v*) can be computed from the other previously defined entities, as follows:

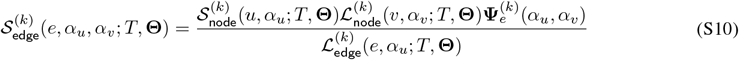

Thus, if all the inside likelihoods, outside likelihoods, and the full likelihood have been computed, we can use Eq. S10 and Eq. S8 to compute all the posterior assignments of states to nodes, in linear-time. With this observation, we can extend the Felsenstein’s prunning algorithm to compute all the inside, outside likelihoods, and posterior assignments, as follows: step (1) compute and store the inside likelihoods at all nodes and branches using Eq. S2 and Eq. S4, in a bottom up tree traversal; step; step (2) compute and store the outside likelihoods at all nodes using Eq. S6, in a topdown tree traversal; step (3) compute the posterior assignments at all nodes using Eq. S10 and Eq. S8, in a tree traversal (either bottom-up or topdown). These values are needed for the E-step of our EM algorithm described later.

### S1.4 The EM algorithm: derivation and pseudocode

#### S1.4.1 Derivation (Proof of Eq. 12 in the main text)

The basic setup of an EM algorithm to optimize *𝒫* (D = **D**; **Θ**) is as follows: in the E-step, compute for every site *k* the posterior probabilities 𝒫 (𝕏^(*k*)^ = **x**|𝔻 ^(*k*)^ = **D**^(*k*)^; **Θ**^*t*^) for all possible realizations **x** of 𝕏 ^(*k*)^. In the M-step, compute **Θ***t*+1 as

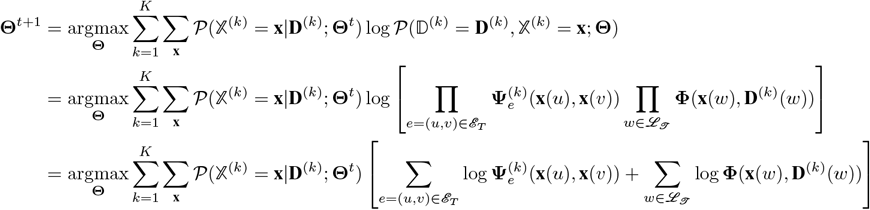

Note that every non-zero entries of **Φ** can only be *ϕ*, 1 − *ϕ*, or 1. Therefore,

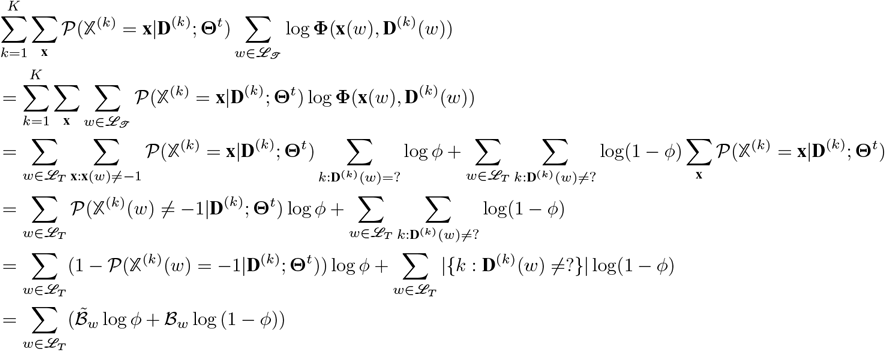

The derivation of the other term is similar: note that all Ψ_*e*_ share the same locations of non-zero entries, and log 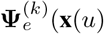, **x**(*v*)) can only be −*δ*_e_ (1 + *ν*), 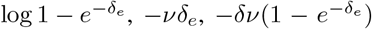, or 0 (plus constants). Therefore, we have (skipping details, as the derivation is very similar to [42]):

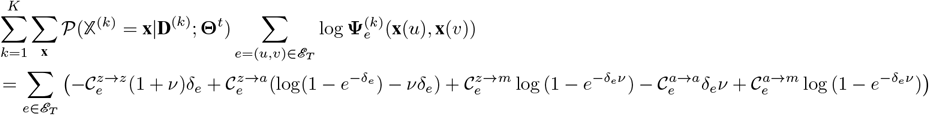

Adding the two terms, we get Eq. 12 in the main text.

### S1.5 Reducing the complexity of the E-step

In the E-step, we need to compute 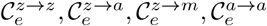, and 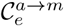 for all branches *e*. Below we introduce an algorithm to compute them in O(*NK*), where *N* and *K* are the number of cells and target sites, respectively. Importantly, the complexity of our algorithm is independent of the alphabet size of any target site. First, we introduce the concepts of *α***-trees**, *α***-clades**, and **z-branches** that will be useful in describing the algorithm.

#### Definition 6.

*Consider a target site k of the lineage tracing data* ***D*** *whose alphabet is 𝒜* ^(*k*)^ *and a character state α* ∈ 𝒜^(*k*)^ \ {0, −1}. *An α****-tree*** *on site k at state α of* ***D*** *is a tree T with leafset ℒ*_*T*_ *such that for all w* ∈ *ℒ*_*T*_, ***D***^(*k*)^(*w*) ∈ {?, *α*}.

#### Definition 7.

*An α****-clade*** *of a tree T with respect to a site k of a lineage tracing data* ***D*** *is a clade of T that forms an α-tree on site k at state α of* ***D***.

Note that the definitions of *α*-tree and *α*-clade include the case where *α* =? (i.e. all leaves of *T* have state “?”). In such a case, we call the tree/clade a *masked tree/clade*.

#### Definition 8.

*A* ***masked-clade*** *of a tree T with respect to a site k of lineage tracing data* ***D*** *is an α-clade of T at k where α* =?. *Any branch belongs to a masked-clade is called a* ***masked branch***. *Any branch that is not a masked branch is referred to as an* ***unmasked*** *branch*.

Note that an *α*-tree where *α≠*? may include some masked clades and masked branches inside it.

#### Definition 9.

*A* ***z-branch*** *of a tree T with respect to a site k of a lineage tracing data* ***D*** *is a branch that does not belong to any α-clade of T on k, for all α* ∈ 𝒜^(*k*)^ \ {0, −1}.

#### S1.5.1 Special properties of *α*-clades and z-branches

##### Theorem 1.

*[az-partition] Consider a target site k of the lineage tracing data* ***D*** *whose alphabet is 𝒜* ^(*k*)^. *Any tree T on* ***D***^(*k*)^ *can be partitioned into* ***edge-disjoint*** *α-clades (each can have a different α* ∈ \{𝒜^(*k*)^ 0, −1} *) and z-branches. This partition is unique and is named the* ***az-partition*** *of T with respect to* ***D***^(*k*)^.

*Proof*. The following is an algorithm to partition any tree *T* into *α*-clades and z-branches.

##### Algorithm 3

Linear-time algorithm to perform az-partition. Input: a tree topology *T* and a target-site **D**^(*k*)^. Output: the az-partition of *T* with respect to **D**^(*k*)^.

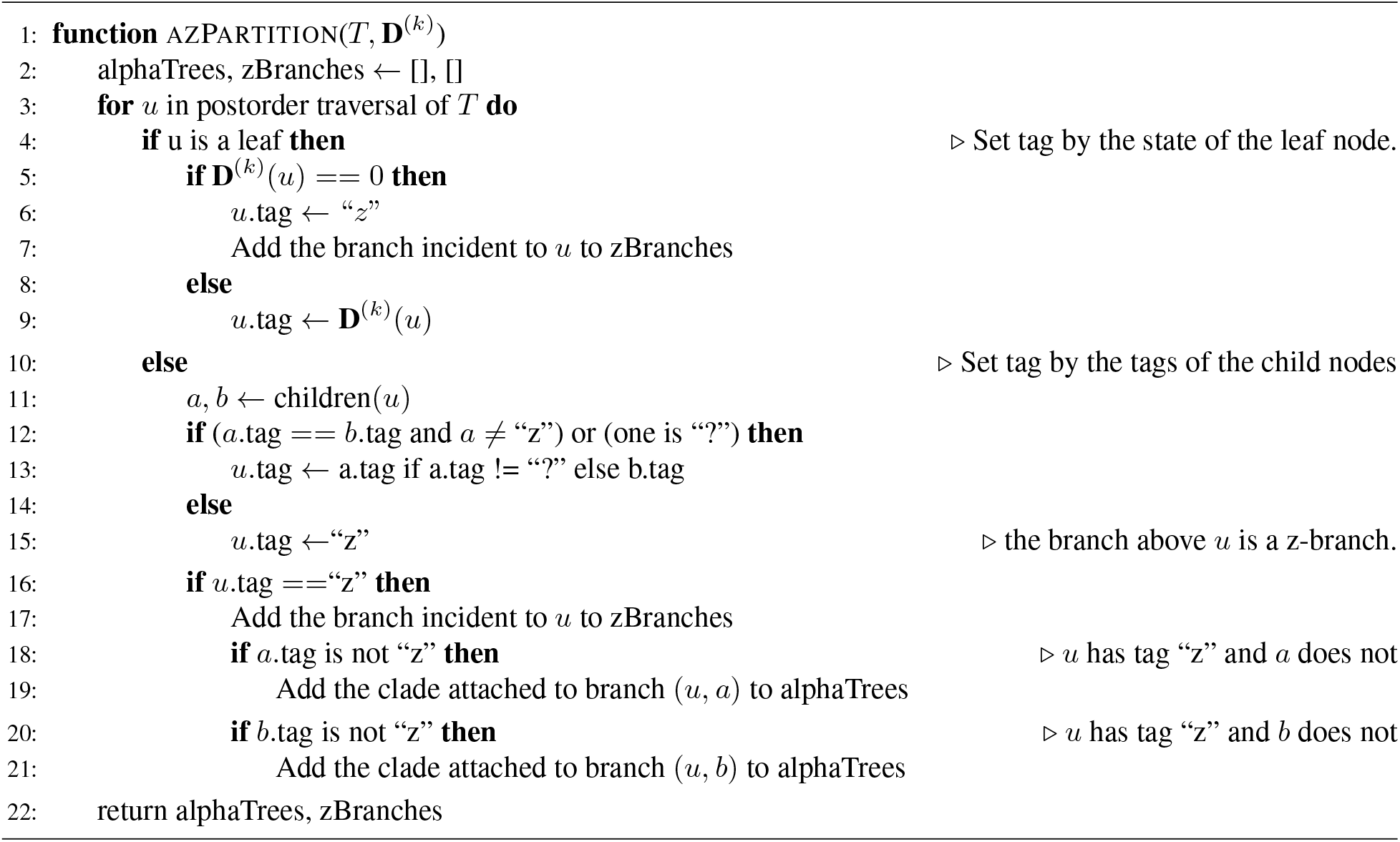

We will analyze this algorithm to demonstrate that it produces a unique az-partition of *T* with respect to *D*^(*i*)^.

The algorithm iterates over every node *u* in the tree *T*, and considers the incident branch, so that the algorithm considers every branch *e*. A branch *e* is either tagged and output as a zBranch (lines 6-7, 15-17), or tagged with the leaf state (lines 12-13). If the branch is tagged with a leaf state *α*, then it must necessarily be output as part of the corresponding *α*-tree at a later iteration. Since a zBranch marks the first branch where not all leaves share the same state, the branches above a zBranch cannot lie in an alphaTree, so alphaTrees must be pairwise disjoint. Therefore, each branch must be output either as part of an alphaTree or as a zBranch, and will only appear once in the output. Thus, we have that the algorithm produces a valid partition.

We should next show that alphaTrees are indeed *α*-clades of *T* for some *α*. For any alphaTree *T*_*α*_ returned by the algorithm, we know that by construction all branches in that alphaTree are tagged with either *α* or “?”, except for the root branch, which is tagged with a *z*. For a branch to be tagged with a leaf state *α*, the incident node *u*’s children have tags in the set {*α*, ? }, recursively down to the leaves (line 11-13) such that all the leaf nodes under this branch must have state in the set {*α*, ? }. Lines 19 and 21, where the entire set of branches below a z-branch is returned, determine that the alphaTrees returned must be a clade. Therefore, by construction we know that the returned trees must be clades that each form an alphaTree with corresponding leafset ∈ {*α*, ? }, as stipulated by Def. 6 and 7.

We should next show that returned zBranches fit Def. 9. zBranches are returned in a few cases: on line 5-6, a zBranch could be returned if the leaf state is 0, which is consistent with Def 9. The other case when a zBranch is returned happens in lines 16-17, where if a node *u* is tagged with a *z*, then the incident branch *b* will be returned as a zBranch. A few scenarios force a node *u* to be tagged with *z*: (1) its two children do not share the same tag, (2) two children are both tagged with *z*. In Case (1), the leaves under branch *b* do not share states ∈ {*α*, ? }, and so this branch cannot belong to any *α*-clade and must necessarily be a zBranch. In Case (2), if both children are tagged with *z*, then the leaves under each child node respectively do not share states ∈ {*α*, ? }∀ *α* ∈ Σ_*i*_ \{ 0,−1}, so that this branch cannot belong to any *α*-clade and must also necessarily be a zBranch. Therefore, all elements of the returned zBranches must be a zBranch, and since all branches are represented at least once, and all alphaTree branches are accounted for, the algorithm must have returned all zBranches.

Putting the above statements together, we want to show that if the algorithm always returns an answer, then the partition must exist. Suppose towards a contradiction the algorithm returns a set of alphaTrees and zBranches, but that no partition exists. However, we saw above that the algorithm necessarily outputs all branches, and each branch appears once as either a zBranch, or as a branch in an alphaTree. Therefore, we must have a valid partition, but we have assumed that no partition exists, and we arrive at a contradiction. We can see that the algorithm must always return an answer given a valid input, since the algorithm output is an assignment for each branch. Consequently, it must be true that if the algorithm returns an answer, then the partition must exist.

Finally, it remains to be shown that the partition is unique. Suppose towards a contradiction that we have two partitions *P*_1_ and *P*_2_ which are both valid partitions for an input tree *T* and data **D**^(*k*)^, but which are different without loss of generality by one branch *b*. Since all branches must show up in both valid partitions, this different branch *b* could be different in tow ways: Case (1): ∈ the set of zBranches in *P*_1_ and ∈ an alphaTree in *P*_2_, or Case (2): ∈ topology alphaTree *T*_*α*_ in *P*_1_ and ∉ topology alphaTree *T*_*β*_ in *P*_2_.

In Case (1), suppose towards a contradiction that it is true a branch *b* could be a zBranch in one valid partition, and in an alphaTree in another valid partition. Then, in valid partition *P*_1_, the leaf states below *b* must not lie in the set {*α*, ?}, and in valid partition *P*_2_, the leaf states do lie in the set {*α*?}. However, both cannot be true, since we have the same tree topology *T* and data **D**^(*k*)^. Therefore, we have arrived at a contradiction, and we know that any branch must either lie in a zBranch or in an alphaTree exclusively, in all valid partitions.

In the other possible Case (2), suppose towards a contradiction that *b* has been assigned to two different alphatrees; that is, *b* is assigned to *T*_*α*_ ∈ *P*_1_ and not assigned to *T*_*β*_ ∈ *P*_2_, where *T*_*α*_ and *T*_*β*_ are topologically the same except for branch *b*. Then, all leaves under branch *b* ∈ *P*_1_ must necessarily lie in the allowed *T*_*α*_ leaf states. In partition *P*_2_, for branch *b* to belong to a different alphaTree, it must be tagged with a different *α*. However, the input data is the same, and so all the leaves below branch *b* must have the same leaf state in both partitions. We have arrived at a contradiction, and thus we have that the partition returned by the algorithm must be unique.

Since missing clades are special cases of alphaTrees, we will not discuss them specially here, except to note that all branches in an alphaTree (masked branches included) will only be output as part of an alphaTree when the ancestral zBranch is processed. We should note that the branch incident to the case where one child has the missing state, and the other child has a nonmissing, nonzero state is not a zBranch.

Thus, we can see that the algorithm must necessarily provide a valid partition of our input tree and data into alphaTrees and a set of zBranches fitting the proposed definitions.

##### Corollary 1.

*Any branch e* = (*u, v*) *of T is either a z-branch, a masked-branch, or an unmasked branch of some α*_0_*-clade of T where α*_0_ ∈ 𝒜^(*k*)^ \ {?, 0, −1}, *with respect to* ***D***^(*k*)^. *In addition, if e is not a z-branch or masked-branch, it has a unique associated α*_0_.

##### Theorem 2.

*[Properties of a z-branch] Given a tree topology T parameterized by* Θ, *a target site k, and a z-branch e* = (*u, v*) *of T on site k. We have the following statements:*

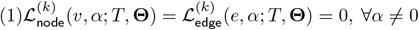

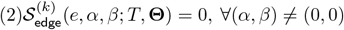

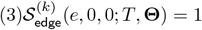

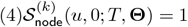

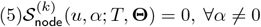

*Proof*. **Proof of Statement (1)**. To prove (1), first we prove that if *e* is a z-branch of site *k*, then leaves in 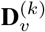 belong to two distinct unmasked states. Suppose towards a contradiction that all leaves in 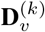 are either in masked state “?” or a same state *α*_0_; by definition of *α*-clade, 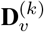 defines an *α*-clade on *T* where *α* = *α*_0_ or *α* =?. By definition, *e* belongs to the clade defined by 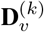, so it is part of an *α*-clade, contradicting the assumption that *e* is a z-branch.

Hence, there exists two leaves *l*_1_ and *l*_2_ under *v* such that 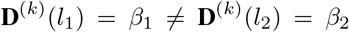, where *β*_1_, *β*_2_ ∈ 𝒜^(*k*)^ \ {?}. Let *w* be the LCA of *l*_1_ and *l*_2_, so 𝔻 ^(*k*)^(*l*_1_) and 𝔻 ^(*k*)^(*l*_2_) are conditionally independent given 𝕏 ^(*k*)^(*w*)).

Recall that 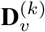 includes **D**^(*k*)^(*l*_1_) and **D**^(*k*)^(*l*_2_). For any *α*≠ 0, we have:

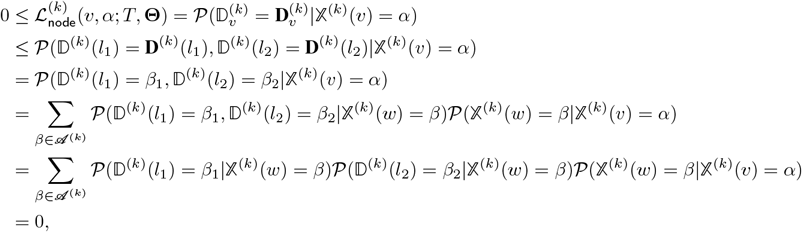

where the last equation holds because all terms inside the last summation are 0: if *β* = 0 or *β ∉* {*α*, −1}, then 𝒫(𝕏 ^(*k*)^(*w*) = *β*|𝕏 ^(*k*)^(*v*) = *α*) = 0; else if *β* = −1 then 𝒫(**D**^(*k*)^(*l*_1_) = *β*_1_|𝕏 ^(*k*)^(*w*) = *β*) = 𝒫 (**D**^(*k*)^(*l*_2_) = *β*_2_|𝕏 ^(*k*)^(*w*) = *β*) = 0 because *β*_1_ ≠? and *β*_2_≠?; otherwise, *β* = *α*, so either 𝒫 (**D**^(*k*)^(*l*_1_) = *β*_1_|𝕏 ^(*k*)^(*w*) = *β*) = 0 or 𝒫(**D**^(*k*)^(*l*_2_) = *β*_2_|𝕏 ^(*k*)^(*w*) = *β*) = 0 because *β*_1_ ≠ *β*_2_ and *α≠* 0. Thus, 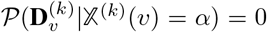 for all *α≠* 0. Now we prove the second equality of (1). For any *α*≠ 0, we have:

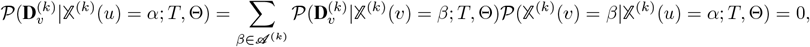

where the second equality holds because 𝒫 (𝕏 ^(*k*)^(*v*) = 0|𝕏 ^(*k*)^(*u*) = *α*; *T*, Θ) = 0 (recall that *α* ≠ 0) and 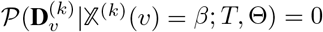 for all *β* ≠ 0.

**Proof of Statement (2)**. Using Eq. S10, we have:

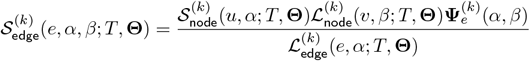

Recall that (*α, β*) ≠ (0, 0), so either *β* ≠ 0 or (*β* = 0 and *α* ≠ 0).

- Case 1: *β* ≠ 0. In this case, 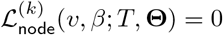 (Statement (1)), so 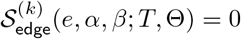.
- Case 2: *β* = 0 and *α* ≠ 0. In this case, 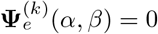 (see Eq. 2 in the main text), so 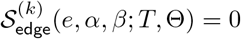.

**Proof of Statement (3)**. It is straightforward to derive Statement (3) from (2):

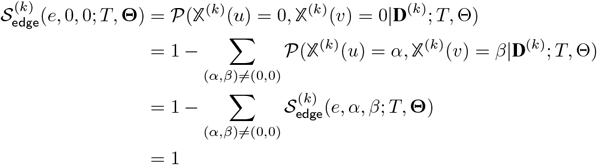

**Proof of Statement (4)**. We prove Statement (4) using Statements (2) and (3):

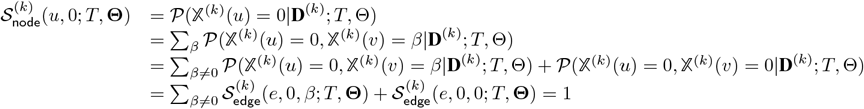

The last equality is justified by Statements (2) and (3).

**Proof of Statement (5)**. For all *α* ≠0, we have:

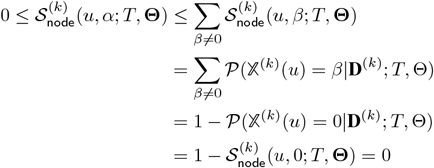

The last equality is justified by Statement (4). Thus, 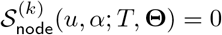 for all *α* ≠0.

##### Theorem 3.

*[Properties of a masked-branch] Consider a masked clade of a tree T at a site k of* ***D*** *and e* = (*u, v*) *is an arbitrary branch of this masked clade. Then we have*

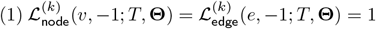

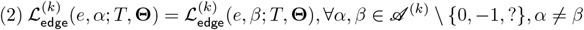

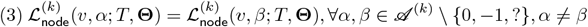

*Proof*. **Proof of (1)**. Let Clade(*v*) denote the leaf set of the clade of *T* below *v*. We have:

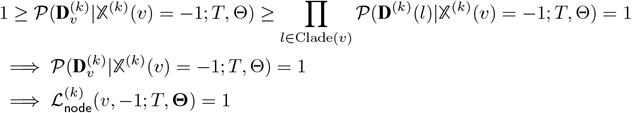

Now we prove the second equality of Statement (1). Indeed:

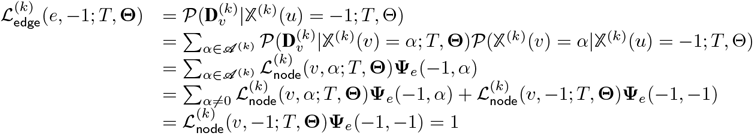

**Proof of (2)**. We will prove by induction. Base case: *e* is a terminal branch (i.e. *v* is a leaf node), so that 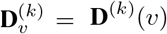. In addition, because *e* is masked, **D**^(*k*)^(*v*) =?. So, we have:

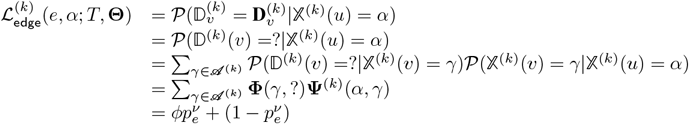

Similarly, we can prove that 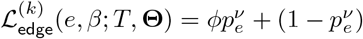, so 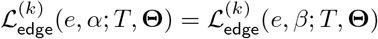.

Now we prove the induction case. If *e* is not a terminal branch, then *v* has two children *v*_1_, *v*_2_. Suppose that the statement is correct for the two branches *e*_1_ = (*v, v*_1_) and *e*_2_ = (*v, v*_2_), we will prove that it is also correct for branch *e* = (*u, v*). For any state *α* ∈ 𝒜^(*k*)^ \ {0, −1, ?}, we have:

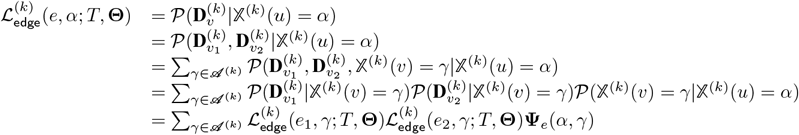

Recall that **Ψ**_*e*_ is a sparse matrix. Because *α* ≠ 0, **Ψ**_*e*_(*α, γ*) = 0 for all *γ* ≠ {−1, *α*}. Therefore, we have:

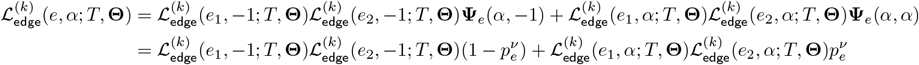

By induction assumptions, for any state *β* ∈ 𝒜^(*k*)^ \ {0, −1, ?} that is distinct from *α*, we have 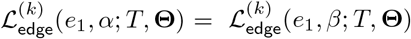 and 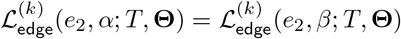 Thus:

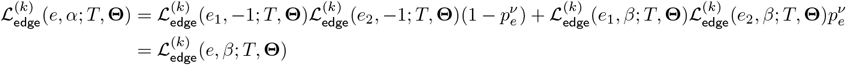

**Proof of (3)**. Similar to (2), we can use induction to prove this statement.

##### Theorem 4.

*[Properties of unmasked branch of an α-tree ] Consider an α-clade of a tree T at a site k of* ***D*** *where α* ≠ ? *and e* = (*u, v*) *is an arbitrary unmasked branch of this clade. Then:*

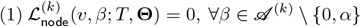

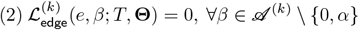

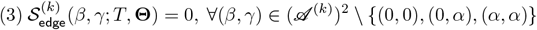

*Proof*. **Proof of (1)** Because *e* = (*u, v*) is an unmasked branch of an *α*-clade of *T* on **D**^(*k*)^, by definition there exists at least one leaf node *l* of *T* under *v* that has **D**^(*k*)^(*l*) = *α* and **D**^(*k*)^(*l*) ∈ 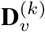. Let *w* be the parent of *l*, and *e*_1_ = (*w, l*). It is easy to see that 𝒫 (𝕏 ^(*k*)^(*l*) = *γ*|𝕏 ^(*k*)^(*v*) = *β*; *T*, **Θ**) ≤ 𝒫 (𝕏 ^(*k*)^(*l*) = *γ*|𝕏 ^(*k*)^(*w*) = *β*; *T*, **Θ**) = **Ψ**_*e*_ (*β, γ*) for all *γ* ∈ 𝒜^(*k*)^. So we have:

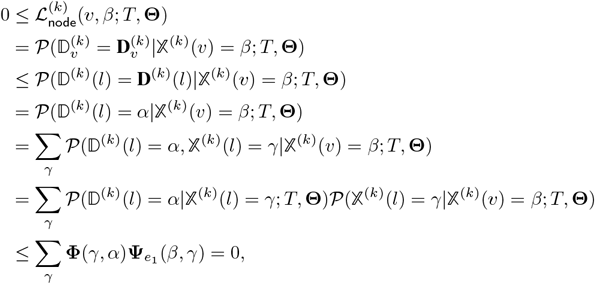

where the last equality holds because of the sparsity of **Φ** and 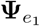 (note that *β* ≠ 0 and *β* ≠ *α*).

**Proof of (2)** Similar to the proof for (1), observe that 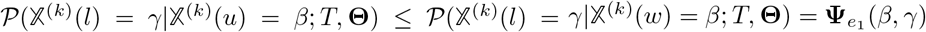 for all *γ* ∈ 𝒜^(*k*)^. So we have:

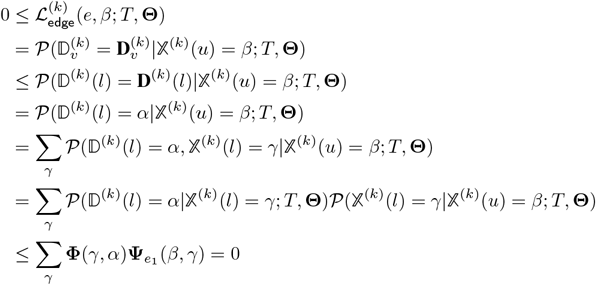

**Proof of (3)** From Eq. S10, for all (*β, γ*) ∈ (𝒜^(*k*)^)^2^ \ {(0, 0), (0, *α*), (*α, α*)}, we have:

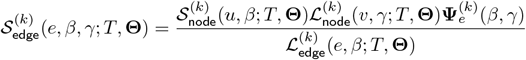

Consider the following exclusive cases:

Case 1: *γ* = 0, then *β* ≠ 0 (because (*β, γ*) ≠ (0, 0)). It is followed that 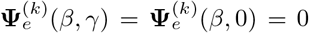, so 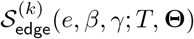.

Case 2: *γ* = *α*, then *β* ≠ *α* (because (*β, γ*) ≠ (*α, α*)). It is also followed that 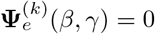, so 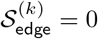
Case 3: *γ* ∉ {0, *α*}. Using Statement (1), we have 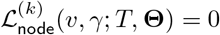, so 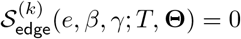.

Thus, 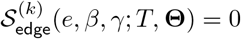 for all (*β, γ*) ∈ (𝒜^(*k*)^)^2^ \ {(0, 0), (0, *α*), (*α, α*)}.

### S1.5.2 The linear-time algorithm

Recall that at iteration *t* + 1 of the EM algorithm, we need to compute the posterior probabilities that are parameterized by **Θ**^*t*^, where **Θ**^*t*^ is the estimate at iteration *t*. Below we describe a new algorithm for the E-step that has lower complexity than that presented in the main text, using the special properties of *α*-clades and z-branches. Because the tree topology *T* and parameter **Θ**^*t*^ are shared for all entities described below, in the rest of this section, we will drop these two parameters in all notations. Let

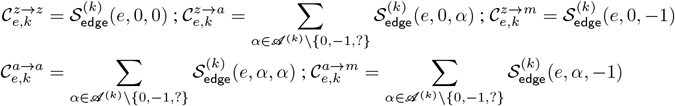

From Eq. 11 (see the main text) and the above equations, it is easy to see that:

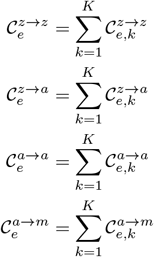

Let 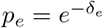. Using theorems 2, 3, and 4, we will prove the followings:

#### Casework for 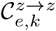

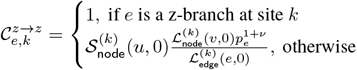

*Proof*. If *e* is a z-branch, Eq. S1.5.2 is a direct corollary of Statement (3) of Theorem 2. Otherwise, from Eq. S10 we have:

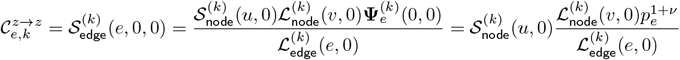

#### Casework for 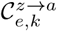

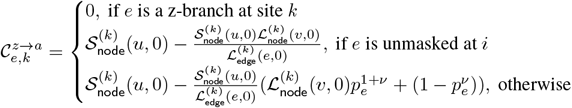

*Proof*. From the definition of 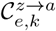 and Eq. S.10, we have:

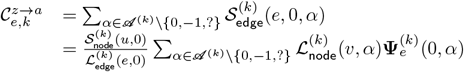

If *e* is a z-branch, then according to Theorem 2, 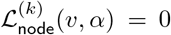 for all *α* ∈ 𝒜^(*k*)^ \ {0, −1, ?}, so 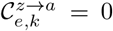. Otherwise, we have:

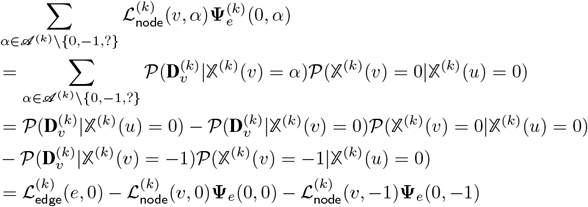

If *e* is a masked branch, then 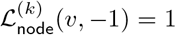; otherwise, *e* is unmasked and is not a z-branch, so 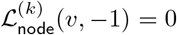 according to Theorem 4. Also, 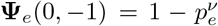 by definition. Thus, from Eq. S1.5.2 and S1.5.2, we get the proposed equation to compute 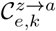.

#### Casework for 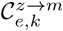

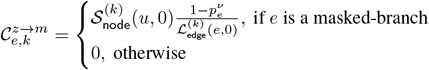

*Proof*. From the definition of 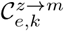 and Eq. S10, we have:

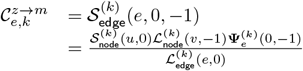

If *e* is not a masked branch, then according to Theorem 2 and Theorem 4, 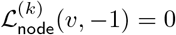, so 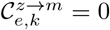. Otherwise, if *e* is a masked branch, then 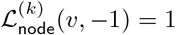 (Theorem 3), and 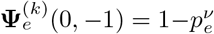. From here we get the proposed equation.

#### Casework for 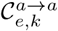

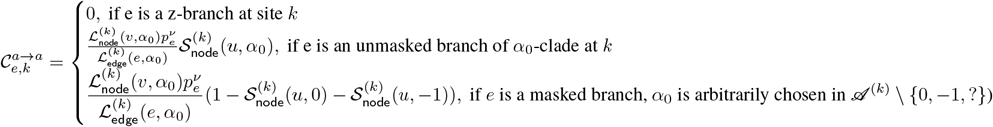

*Proof*.

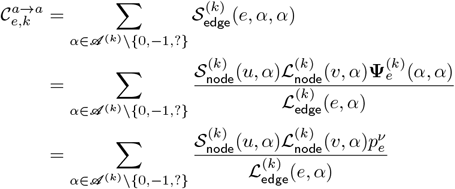

Case 1: If *e* is a z-branch, then 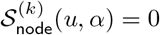 for all *α* ∈ 𝒜^(*k*)^ \ {0, −1, ?} (Theorem 2), so 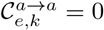.

Case 2: If *e* is an unmasked branch of some *α*_0_-clade, then 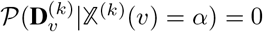 for all *α* ∈ 𝒜^(*k*)^ \{ 0, −1, ?, *α*_0_} (Theorem 4), so the summation is reduced to the only term containing *α*_0_. Thus:

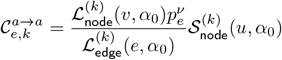

Case 3: If *e* is a masked branch, then from Statements (2) and (3) of Theorem 3, we have:

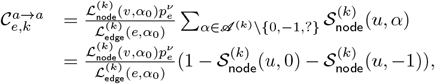

where *α*_0_ is arbitrarily chosen in 𝒜^(*k*)^ \ {0, −1, ?}

#### Casework for 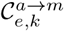

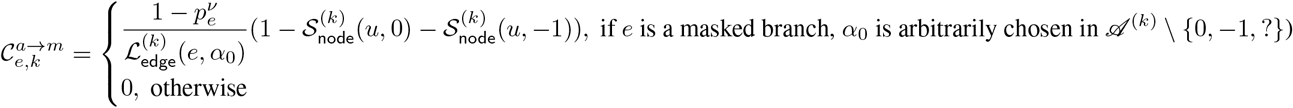

if *e* is a masked branch, α_0_ is arbitrarily chosen in 𝒜^(*k*)^ \{0, −1, ?}

*Proof*. From the definition of 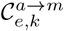 and Eq. S10, we have:

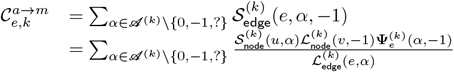

If *e* is not a masked branch, then according to Theorem 2 and Theorem 4, 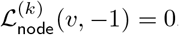, so 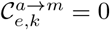. Otherwise, if *e* is a masked branch, then 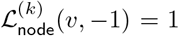 (Theorem 3), 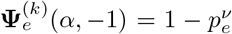 for all *α* ∈ 𝒜^(*k*)^ \ {0, −1, ?}. Combining with Statements (2) and (3) of Theorem 3, we have:

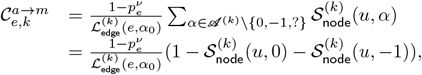

where *α*_0_ is arbitrarily chosen in 𝒜^(*k*)^ \ {0, −1, ?}

Thus, for any branch *e* = (*u, v*), if *e* is a z-branch of site *k*, then each of the 5 entities 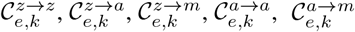 is either 0 or 1 (Theorem 2). Otherwise, to compute them we need the following 6 entities:

- 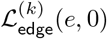
- 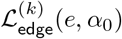
- 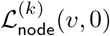
- 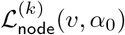
- 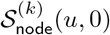
- 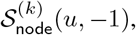

where *α*_0_ is arbitrarily chosen in 𝒜^(*k*)^ \ {0, −1} if *e* is a masked-branch; otherwise, *e* belongs to an *α*-clade where *α* is uniquely defined and we let *α*_0_ = *α*.

#### Computing 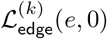 and 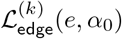

Base case: if *v* is a leaf node, then

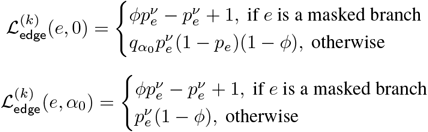

Recursion: if *v* has two children, *v*_1_ and *v*_2_, then let *e*_1_ = (*v, v*_1_), *e*_2_ = (*v, v*_2_), we have:

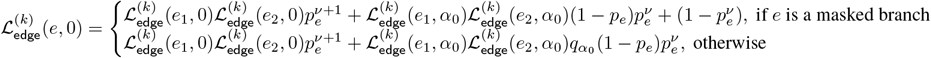

#### Computing 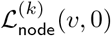 and 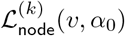

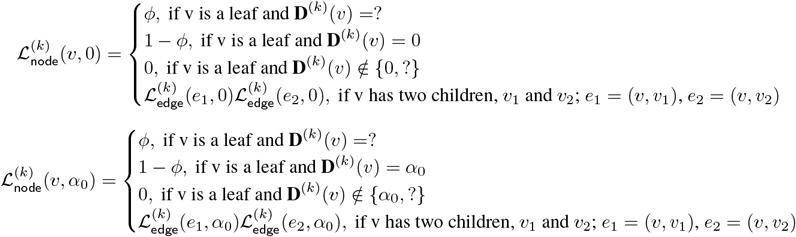

#### Computing 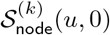 and 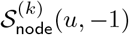

Consider a branch (*u, v*). From Eq. (S8), we have: if *u* is the root, Then 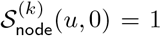 and 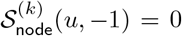. Otherwise, to compute these two entities we need 5 other entities: (1) 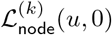 (2) 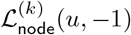, (3) 𝒫 (**D**^*(k)*^), (4) 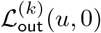, and (5) 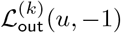. Among them, (1) and (2) are already computed (see the above section). Entity (3) is simply the full likelihood of site *k*, so it is equal to 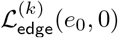 where *e*_0_ is the root branch. To compute (4) and (5) (i.e. the outside likelihoods), we can use Eq. (S6), which gives a top-down recursive formula. Let *u*_0_ be the parent of *u, e* = (*u*_0_, *u*) be the branch connecting *u*_0_ and *u*, and *u*_1_ be the sister of *u* (if there exists one) and *e*_1_ = (*u*_0_, *u*_1_). From Eq. (S6), it is straightforward to derive the following:

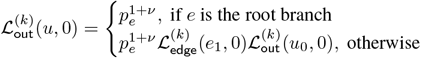

Thus, we have a top-down recursion to compute 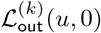. The computation of 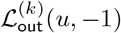, however, is more complicated. A naive application of Eq. (S6) requires the outside likelihoods with respect to all states *β* at the parent of *u*. Therefore, it takes 𝒪 (*M*) time to compute 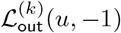for each node *u*, so the overall complexity is 𝒪 (*NKM*) where *M* is the maximum alphabet size. Nevertheless, we can still employ the special properties of the model to reduce the algorithm to 𝒪 (*NK*), as shown below.

Let *v* be the parent of *u, e* = (*v, u*) ∈ *ℰ*_*T*_ be the branch connecting *v* and *u, w* be the sister of *u*, and *e*_1_ = (*v, w*). Using Eq. S6, it is straight forward to prove the followings:

- If *e*_1_ is a z-branch, then 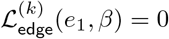 for all *β≠* 0. Therefore, 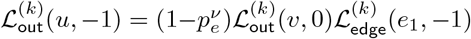
- If *e*_1_ is an unmasked branch of an *α*_0_-clade, then 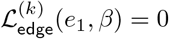 for all *β ∉* {0, *α*_0_}. Therefore, 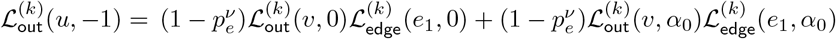
- If *e*_1_ is a masked branch, then 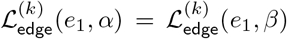 for all *α ≠β, α, β* ∈ 𝒜^(*k*)^ \ {0, −1, ?} (The-oreom 3). Let *α*_0_ be an arbitrarily chosen character state in 𝒜^(*k*)^ \ {0, −1, ?}. Then 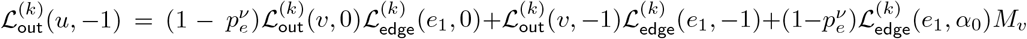, where 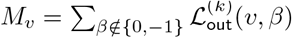.

From the above equations, we see that in all cases 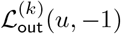 can be computed from 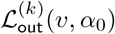, *M*_*v*_, and other known entities. Therefore, what left to be done is deriving top-down recursive formulas for 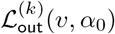 and *M*_*v*_. Below we show the formulas and leave the derivation to the reader.

#### Recursive formula for *M*_*v*_

Consider a branch *e* = (*u, v*), let *w* be the sister of *v* and *e*_1_ = (*u, w*). Recall that by definition, 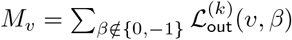. Below is the topdown recursive formula to compute *M*_*v*_ from *M*_*u*_ and other known entities:

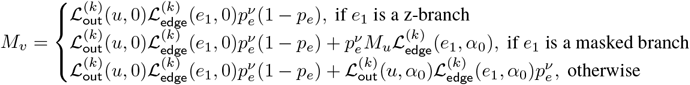

#### Recursive formula for 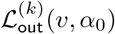

Consider a branch *e* = (*u, v*), let *w* be the sister of *v* and *e*_1_ = (*u, w*). For any *α*_0_ ∉ {0, −1}, we have the following topdown recursive formula to compute 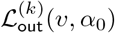 from 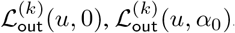, and other known entities:

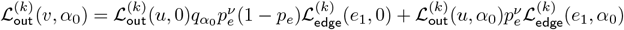

### S1.6 Solving the M-step

#### S1.6.1 Proof of convexity by variable blocks

Recall that at iteration *t* of the proposed EM algorithm, in the M-step we need to solve the following optimization problem:

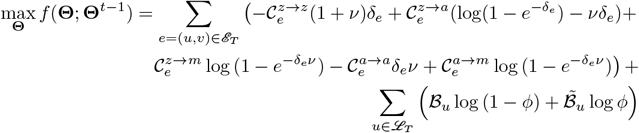

Below we prove that *f* (**Θ**; **Θ**^*t*−1^) is concave with respect to {*δ*_*e*_}, *ν*, and *ϕ* separately. Note that all variables are separable, so to prove concavity, we need to show that the second-order partial derivatives are negative. Indeed, we have:

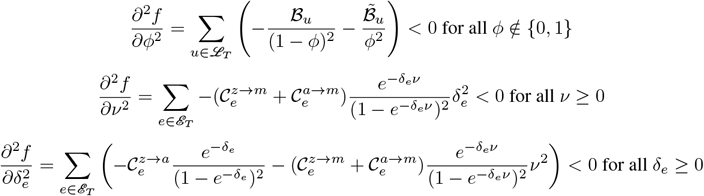

#### S1.6.2 The block coordinate ascent algorithm

We initialize **Θ**^*t*^ ← **Θ**^*t*−1^, then successively minimize *f* along the coordinate of *ϕ, ν*, and the block of *{δ*_*e*_*}* while fixing the others, and iterate until convergence. In other words, let *ϕ*^(*t*,1)^ = *ϕ*^(*t*)^, *ν*^(*t*,1)^ = *ν*^(*t*)^, and *{δ}*^(*t*,1)^ = *{δ}*^(*t*)^; at each iteration p of coordinate ascent, we find *ϕ*^(*t,p*+1)^, *ν*^(*t,p*+1)^, and *{δ}*^(*t,p*+1)^ such that:

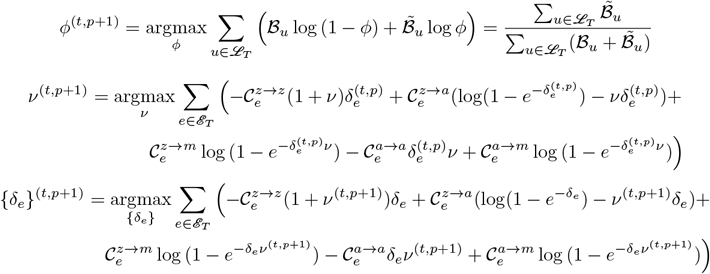

While *ϕ*^(*t,p*+1)^ has a closed-form, *ν*^(*t,p*+1)^ and *{δ*_*e*_*}* ^(*t,p*+1)^ require solving convex optimization problems. We use the CVXPY package [12, 1] with the MOSEK solver [3] to solve them.

### S1.7 Simulated annealing for topology search

We repeat the following procedure, starting with iteration *t* = 1, until convergence or *t > t*_max_:

1. At iteration *t*, do the following steps:
  a. Propose a new topology *T* ^(*t*)^ using an NNI move on *T* ^(*t*−1)^.
  b. Optimize for **Θ**^(*t*)^ using EM algorithm
  c. Compute the log-likelihood *l*^(*t*)^ = log *L*(*T* ^(*t*)^, **Θ**^(*i*)^; **D**)
  d. Go to (2)
2. If *l*^(*t*)^ *> l*^(*t*−1)^, then accept *T* ^(*t*)^ and **Θ**^(*t*)^ and go to (3). Otherwise, compute the temperature Temp(t) and acceptance probability *p*(*t*), as follows:

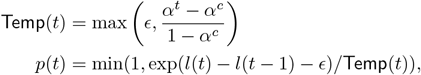

where *c, α* and *ϵ* are hyperparameters controlling the simulated annealing procedure. By default, we set *ϵ* = 1*e* − 12, *c* = 20, and *α* = 0.9. Accept *T* ^(*t*)^ and **Θ**^(*t*)^ with probability *p*(*t*) and go to (3).
3. If *T* ^(*t*)^ and **Θ**^(*t*)^ are accepted in (2), then increase *t* and iterate back to step (1). Otherwise, reject *T* ^(*t*)^ and **Θ**^(*t*)^; without increasing *i*, turn back to step (1) and try another NNI move of *T* ^(*t*−1)^. If there is no NNI move left or the likelihood improvement is below a predefined threshold *η*, stop the procedure and return *T* ^(*t*−1)^ and **Θ**^(*t*−1)^.

## S2 Supplementary data Contents

### S2.1 Evaluation metrics

#### S2.1.1 The Robinson-Foulds (RF) error

For simulated data and the intMemoir dataset where the ground-truth topology is known, we measure topological error by the *RF error*.

The RF distance between two trees *T*_1_ and *T*_2_ on a same leafset is the sum of false positives (FP - bipartitions in *T*_2_ not present in *T*_1_) and false negatives (FN - bipartitions in *T*_1_ not present in *T*_2_). We normalize the RF distance by the total number of internal edges in both trees. Thus:

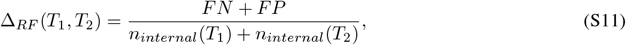

where *n*_*internal*_ denotes the number of internal edges of a tree. The RF error of an estimated topology is simply its normalized RF distance to the true topology. We use *root-mean-square error (RMSE)* to compute the error in estimating *ϕ* and *ν*. Specifically, for each model condition where *θ*^*^ = (*ϕ*^*^, *ν*^*^) is the true set of parameters, the error of an estimate 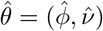 is 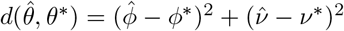. The RMSE of the estimates across *m* replicates is

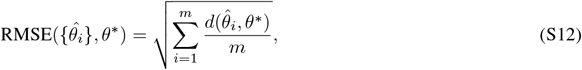

where *m* = 50 in all model conditions.

#### S2.1.2 Migration Cost

For the sample 3724 NT T1 All of KP-tracer data where topology is re-estimated using LAML, we compare the trees in terms of *migration cost*, which is the phylogeny’s parsimony cost on the annotated anatomical labels of the cells. This metric is the same as that used in [39], and was introduced and implemented in MACHINA [14]. Below we briefly describe the construction of a *migration graph*; see [39] and [14] for more details.

1. First, infer the ancestral labeling using MP (i.e. migration annotations on the internal nodes).
2. Collapse all branches that share identical labels between the parent node and child node.
3. Group all unique (parent state, child state) edges. Each state tuple should be annotated with the number of times edges labeled with (parent state, child state) were observed in the tree. These correspond to migration events.

The *migration cost* is the sum of the edge weights.

#### S2.1.3 Variations on the Migration Cost

The KP-Tracer sample (`3724_NT_ALL`) includes two useful meta annotations on the observed cells. First, we have the anatomical site labeling (e.g. primary tumor, lung metastasis 1, lung metastasis 2, lung metastasis 3, soft tissue metastasis). This indicates where in the body each cell was sampled from. Second, we have a “refinement” of this anatomical site labeling for certain tumors, that indicates the approximate spatial location of the sample in that tumor (e.g. primary tumor can be broken into 15 components).

As defined above, a migration graph is defined by the tree and the clustering. Thus, given a tree topology *T* and a clustering *SC* for spatial clustering and *AC* for anatomical clustering, we can get migration graph *T* + *SC* → *MG*_*SC*_ and *T* + *AC* → *MG*_*AC*_. We presented results on the anatomical clustering in the main body of the paper. We include additional results on *MG*_*SC*_, and compute different costs on this migration graph (below) in the Supplementary Results section for KP-Tracer S2.7.

1. Total Cost: The migration cost of *MG*_*SC*_(*T*_*LAML*_) and *MG*_*SC*_(*T*_*CassH*_).
2. Induced Anatomical Cost: Using the parsimonious ancestral labeling inferred using the spatial clustering *SC*, infer the number of transitions between anatomical sites.
3. Spatially-aware Migration Cost: Within the **primary tumor only**, each transition is weighted as follows: migration cost * length of path between spatial locations.
4. Constrained Parsimony: Given a tree topology *T*, we first compute a parsimonious ancestral labeling solution given the anatomical clustering *AC*. Then, for clades within each anatomical site, we compute a parsimonious ancestral labeling solution given the spatial clustering *SC*.

#### S2.1.4 Mutual information with gene expression

For the 6 samples of KP-tracer where branch lengths are evaluated, we use gene expression data to benchmark the branch lengths estimated by MP and ML. We construct two pairwise *phylogenetic distance matrices*, one for MP and one for ML, based on the corresponding estimated branch lengths. In addition, we add another distance matrix computed solely from tree topology (i.e. setting all branch lengths to 1), used as control. We preprocess the gene expression data by performing dimensionality reduction, using PCA (10 PCs), scVI (10 PCs), and UMAP (2 PCs). We use the published vectors provided in KP-tracer for scVI and UMAP, and perform PCA using SCANPY [51]. We constructed 3 pairwise *gene expression distance matrices* corresponding to each of those 3 dimensionality reduction techniques, using Euclidean distances. Thus, we have 6 distance matrices for each KP-tracer sample: 3 constructed from phylogenetic distances and 3 constructed from gene expression distances. We pair each phylogenetic with each gene expression distance matrices, producing 9 phylo-expression pairs. For each of these 9 pairs, we compute the non-parametric mutual information [27, 36] using scikit-learn [34]. These mutual information scores are used to benchmark MP, ML, and topological distances.

### S2.2 Simulation procedure

We simulate a model tree of 1024 leaves (i.e. 10 cell generations) using the birth-only model (implemented in Cas-siopeia). Branch lengths in time unit are drawn according to a lognormal distribution with mean 1 and standard deviation 0.1. We simulate sequences of length 30 under our generative model, with hyperparameters *q*^(*k*)^ at each site *k* drawn from that of the trunk-like single-cell lineage tracing mouse (TLSCL) dataset (unpublished data). We also match two other statistics of the TLSCL: the expected proportion of non-zero entries (64%) and missing data (25%). To achieve an expected proportion of 64% non-zeroes, we use a molecular clock that has *constant mutation rate λ* = 0.095. To obtain a 25% overall missing proportion, we calibrated *ϕ* and *ν*for each of the 5 model conditions, as summarized in Table S2. Each simulated character matrix is subsampled to 250 to reflect sparse sampling, and the model tree is pruned to the same subset. We repeat the subsampling procedure 5 times for each character matrix, creating 250 replicates in total.

**Table S2:** Simulated dataset under five model conditions. All model conditions was simulated with 25% missing data; the attribution of the missing entries to heritable missing versus dropout depending on the model condition.

| Model condition | Heritable percent | Dropout percent | Heritable rate ($\nu$) | Dropout rate ($\phi$) |
| --- | --- | --- | --- | --- |
| h0d100 | 0% | 100% | 0 | 0.25 |
| h25d75 | 25% | 75% | 0.065 | 0.2 |
| h50d50 | 50% | 50% | 0.134 | 0.143 |
| h75d25 | 75% | 25% | 0.208 | 0.077 |
| h100d0 | 100% | 0% | 0.288 | 0 |

#### S2.2.1 Benchmarking LAML

Using this simulated dataset, we benchmark LAML against other methods: Greedy method (based on the principle of perfect phylogeny, used as the baseline), Startle [39], Neighbor Joining [38] (implemented in Cassiopeia, with weighted Hamming distances [25]), and DCLEAR. For the Greedy method, we use the implementation of Cassiopeia [25]). We run Startle in NNI mode with 250 iterations (Startle-NNI) and use the topology estimated by Greedy method as the starting tree. LAML was run using Startle-NNI as the starting tree; simulated annealing NNI with EM optimization was performed until convergence of likelihood.

We used the following commands to run each method. For all Cassiopeia-implemented methods, we used the wrapper script: `cassiopeia_solvers_with_pickle.py`. Following the default tutorial pipeline, we allow Cassiopeia to collapse all mutationless edges, producing trees with polytomies.

1. Startle-NNI: `python startle.py <seed_tree> <character_matrix> -e <mutation_priors> --iterations 250 --output tree.nwk`
2. Greedy method (Cassiopeia-Greedy): `cas.solver.VanillaGreedySolver()`
3. Neighbor Joining (Cassiopeia): `cas.solver.NeighborJoiningSolver( dissimilarity_function=cas.solver.dissimilarity.weighted_hamming_distance, add_root=True)`
4. LAML: `python run_problin.py -t <starting_tree> -c <character_matrix> -p <mutation_prior> -o <output> -v --delimiter comma --nInitials 1 --topology_search --ultrametric > <logfile> 2>&1`

To run the Cassiopeia-Hybrid (CH) and Cassiopeia-ILP (CILP) versions, we tried Cassiopeia Release (tag 2.0.0) https://github.com/YosefLab/Cassiopeia/releases/tag/2.0.0, as well as the development version (last updated Aug 8 2023, commit hash: `f89530145adad929fc47a2e80f70a93c6af2cf3f`). Unfortunately, we were unable to run either of these methods on our simulated dataset. CH failed on all of the simulated data inputs by throwing an index error within the Cassiopeia code. CILP failed on approximately half the simulated data inputs (some with Gurobi errors and some with index errors from within the Cassiopeia code).

There are two ways of running DCLEAR. The first, DCLEAR using a *k*-mer based approach, is referred to as `DCLEAR (KRD)`, and the second, DCLEAR using a training-based approach, is referred to as `DCLEAR (WHD)`. Here we report results for `DCLEAR (KRD)` only, because `DCLEAR (WHD)` produced segmentation fault errors. We followed the tutorial available here: https://ikwak2.github.io/tmphtml/Example_subchallenge2. To run DCLEAR (KRD), we note that `DCLEAR` assumes characters across different sites share an alphabet. However, when we converted our simulated data to follow this convention, `DCLEAR` could not scale to handle alphabet sizes of approximately 250. Instead, we report results from running `DCLEAR` directly on the input character matrices, which were also directly provided to the other benchmarked methods. We followed this `DCLEAR (KRD)` vignette: https://colab.research.google.com/gist/gongx030/653a76bffc4ee6ff41499e0026b6d39a/krd.ipynb.

Given a fixed topology, we used LAML to estimate numeric parameters (branch lengths, missing data rates). We used the following command to do so: `python run_problin.py -t <fixed_topology> -c <character_matrix> -p <mutation_priors> -o <output> -v --delimiter comma --nInitials 1 --ultrametric`

#### S2.2.2 Leveraging the Statistical Model in Distance-Based Approaches

One of our methodological contributions in this paper is a new probabilistic model of CRISPR/Cas9 lineage tracing. In this section we elaborate on the applications of this new statistical model.

Using the statistical model, we can estimate maximum likelihood distances between any two sequences. To explore how maximum likelihood distances under our statistical model compare to other ways of computing distances, we compute a distance matrix using several different approaches and run the widely-used distance-based method Neighbor Joining (NJ). This comparison is run on simulated data under all model conditions (varying missing data type composition).

We explain each pipeline in greater detail below:

1. wHD (weighted Hamming Distance): Standard Hamming distance is modified by provided weights. These weights are negative log transformed from the prior transition probabilities for each site’s alphabet. wHD is recommended within the Cassiopeia implementation of NJ [**?**, 25]. `cas.solver.NeighborJoiningSolver(dissimilarity_function= cas.solver.dissimilarity.weighted_hamming_distance, add_root=True)`
2. modified-AC (modified Allelic Coupling distance): Standard allelic coupling distance [52] is modified here by consideration of the missing state. This distance matrix is passed to PHYLIP-NJ to construct a tree for each sample. We use the `get_wAC.py` script to calculate the distance matrix.
3. MLpair: Given an input character matrix, we use LAML to estimate maximum likelihood distances for each rooted pair of leaves. After building this distance matrix, we run PHYLIP-NJ to construct a tree for each sample. We use the `split_pairwise.py` script to split the input distance matrix into pairs, and compute the maximum likelihood distances for each rooted pair.
4. MLall: Running LAML in the default way, we estimate maximum likelihood distances over all pairs of leaves at once, exploring tree topology space to find a maximum likelihood tree.

### S2.3 The KP-tracer data

We benchmark LAML on KP-tracer, a recently published dynamic lineage tracing experiment in mouse models of metastatic lung adenocarcinoma [52]. The data come with character matrices for multiple samples, each has its own per-site alphabet hyperparameters **q**^(*k*)^.

To evaluate tree topology, we use the largest tumor dataset, `3724_NT_T1_All`, which consists of 21, 108 cells and 9 target sites. Originally, Yang et al. published a phylogeny using the Cassiopeia-Hybrid (Cass-hybrid) method [52]. Later, Sashittal et al. [39] published another phylogeny using Startle-NNI that yielded a more parsimonious explanation of the metastatic process compared to the published Cass-hybrid phylogeny. We use the migration cost metric, which we discuss below Supplementary Section S2.1.2. To run LAML, we first remove cells with identical sequences from the original character matrix, resulting in 1, 461 unique sequences. Then we prune the published Startle-NNI tree to this set of 1, 461 sequences and use it as starting tree. We search for the tree that optimizes the log-likelihood of the sequence data using the provided hyperparameters. We run topology search in LAML for 450 iterations to optimize for log-likelihood, which is improved from -12288 in the Startle-NNI tree to -12208 in the LAML tree. Finally, we add the duplicated sequences to the tree as polytomies.

To evaluate branch length estimation, we use 6 other samples of KP-tracer that have a medium-high number of target sites (29 to 73) and moderate number cells (29 to 294); these are: 3432_NT_T1, 3435_NT_T4, 3520_NT_T1, 3703_NT_T1, 3703_NT_T2, and 3703_NT_T3. For these 6 selected samples, we use the published tree topologies (estimated by Cassiopeia) and estimate branch lengths using LAML (here after referred to as dML), and compare our estimate to that of maximum parsimony (here after referred to as dMP).

### S2.4 The intMEMOIR data

The MEMOIR [17] technology encodes DNA barcodes, and performs lineage tracing by accumulating variable edits in the encoded synthetic target array, and which allows readout by imaging. The intMEMOIR technology [10] permits extended recording and germline transmission, so that the true evolutionary history of sampled cells is recorded. In the intMEMOIR technology, only two edits are possible (either deleting or inverting the target region), so that only three states are possible at any of the sites. We should note this is in sharp contrast to the technology used in [7], where each site can have a unique, variably-sized alphabet of possible edits. In this dataset, we have 106 samples, each with 10 sites, with a maximum of 40 cells and a minimum of 4 cells. The intMEMOIR dataset was used in the original DREAM challenge [20], making it a widely used benchmarking dataset. TiDeTree [40], which takes a Bayesian approach to inferring phylogenies, and was intended for use as a module in the BEAST package, was tested on the intMEMOIR dataset.

The TiDeTree [40] paper details that the topologies for all 106 intMEMOIR samples were jointly estimated, with shared scarring rates and population dynamic parameters across all trees, and assuming a molecular clock rate. They benchmark against AMbeRland, Guna lab, Cassiopeia, Jasper06, pRennert and RnlLabs. The author kindly made their resulting maximum clade credibility (MCC) trees available to us. We post-processed these trees as TiDeTree’s supplement specified, by running BEAST’s TreeAnnotator tool (with the “maxium clade credibility” option to create a summary tree.

We benchmarked LAML against Startle-NNI, Cassiopeia and TiDeTree using the following commands:

1. Startle-NNI: `python startle.py $seed_tree $character_matrix -e $mut_priors --iterations ${iters} --output ${output_dir}/startlenni_tree_collapsed.newick`
2. Cassiopeia: We ran Cassiopeia-Hybrid with a cut-off of 100 cells, so that we effectively report results for Cassiopeia-ILP on all intMEMOIR samples. cas.solver.HybridSolver(cas.solver.VanillaGreedySolver(), cas.solver. ILPSolver(convergence_time_limit=100, convergence_iteration_limit=1, maximum_potential_graph_layer_size=1000), cell_cutoff=100)}
3. LAML: python /n/fs/ragr-research/projects/problin/run_problin.py -t $tree -c $msa -p $prior -o $prefix.txt -v --delimiter comma --nInitials 1 --topology_search --ultrametric --maxIters 2500 --parallel > $prefix.log 2>&1
4. TiDeTree: We adapted the provided example scripts https://github.com/seidels/tidetree/blob/main/examples/, using the default parameters and provided the sequence information. We installed BEAST 2.7 and used the following recommended compiled jar file on the adapted example XML file: `java -jar bin/tidetree.jar examples/adapted_example.xml`

### S2.5 Supplementary Results for Simulated Data

**Table S3:** Robinson-Foulds error (RF), weighted maximum parsimony (WMP), and log likelihood (LLH) of the methods on simulated data, summarized over all five model conditions (250 replicates). For each pair of method and metric, the percentage of times the method scored the best according to that metric is shown.

|  | **Greedy method** | **Startle-NNI** | **LAML** |
| --- | --- | --- | --- |
| RF distance to the true tree | 0% | 0% | **100%** |
| Weighted parsimony cost | 0% | **84%** | 16% |
| Log-likelihood | 0% | 0% | **100%** |

### S2.6 Benchmarking LAML

**Figure S1:**
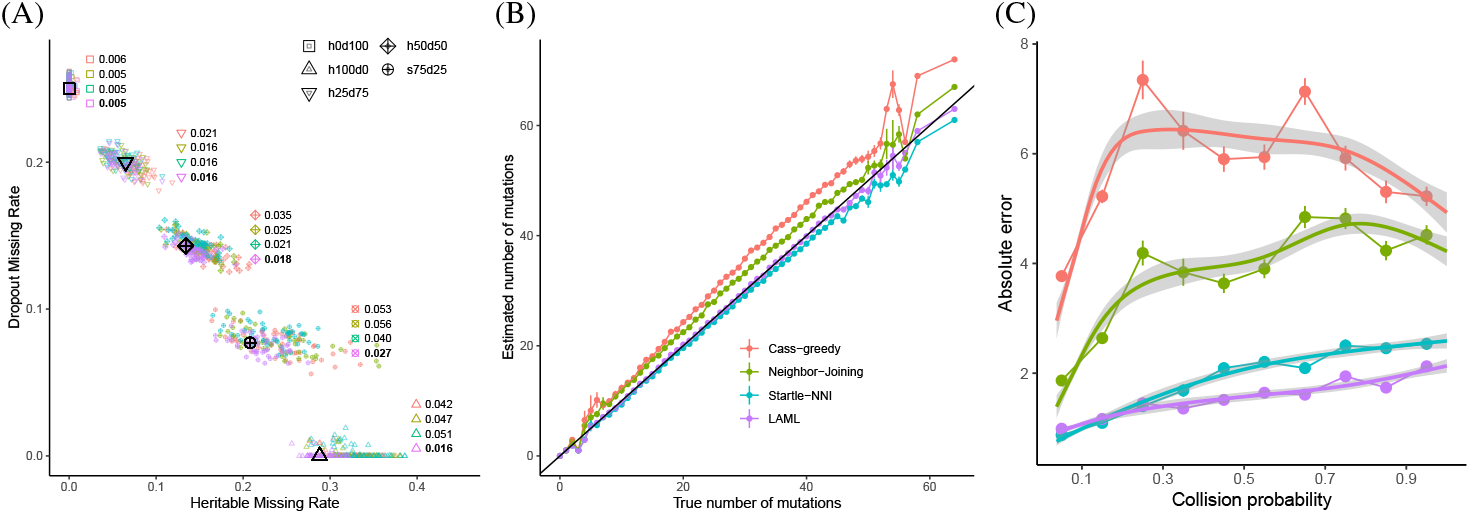
Supplementary results on the simulated data. (A) Estimation of *ϕ* and *ν*. True values of each model condition are shown in black, while the estimates using different tree topologies are shown in different colored, together with the root-mean-square-error (RMSE) of the estimates on each model condition. (B) Estimated versus true number of mutations. Each dot is the average estimated value by each method around one true value, shown with error bar. (C) Absolute error in estimating number of mutations versus collision probability of each target site. Collision probability (x-axis) is computed for each target site across all model condition. The x-axis is discretized into 10 bins and absolute error is averaged for each bin, shown with error bar. For (B) and (C), results are combined for all model conditions.

In Figure S2, we have simulated data results illustrating that LAML produces the most likely tree, as well as relatively parsimonious trees. However, we should note that the most parsimonious tree (Startle-NNI) shows very different RF error (2). Thus, likelihood and RF are more correlated than parsimony and RF are, suggesting that likelihood may be a better metric for scoring trees.

**Figure S2:**
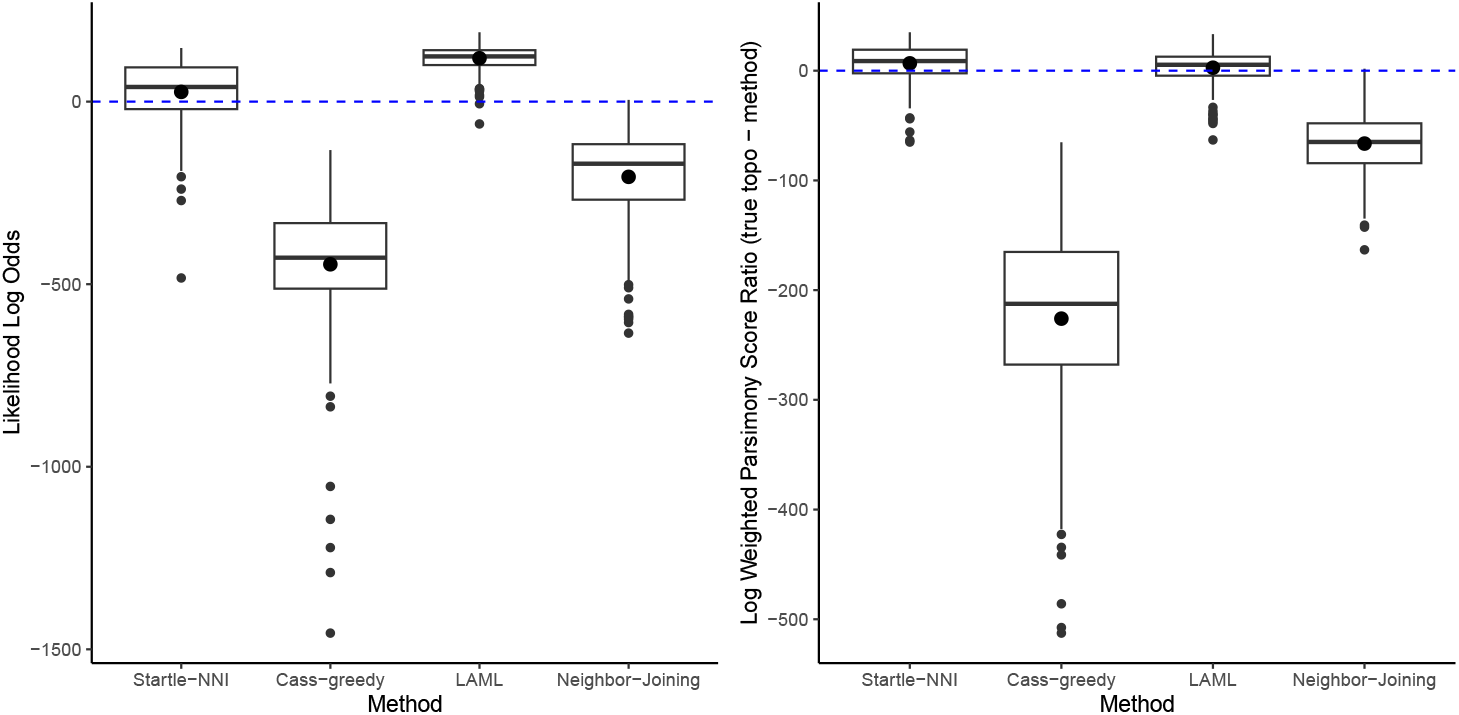
Comparison of the benchmarked methods based on two metrics: estimated log likelihood (calculated by LAML) to the true tree likelihood on simulated data (left); and weighted parsimony score (right).

In Figure S3, we show an additional comparison to Cassiopeia-ILP. Unfortunately, Cassiopeia-ILP was unable to run on all of the simulated data, finishing on 44*/*250 samples (h0d100: 13, h100d0: 8, h25d75: 15, h50d50: 4, h75d25: 4). Of those that finished with an error, 100 of the jobs did not finish running given 168 hours of runtime (7 days) and, 48 jobs ran out of memory (given 4G per job), 26 finished with a GurobiError, and 32 finished with an “IndexError” from within the Cassiopeia code.

**Figure S3:**
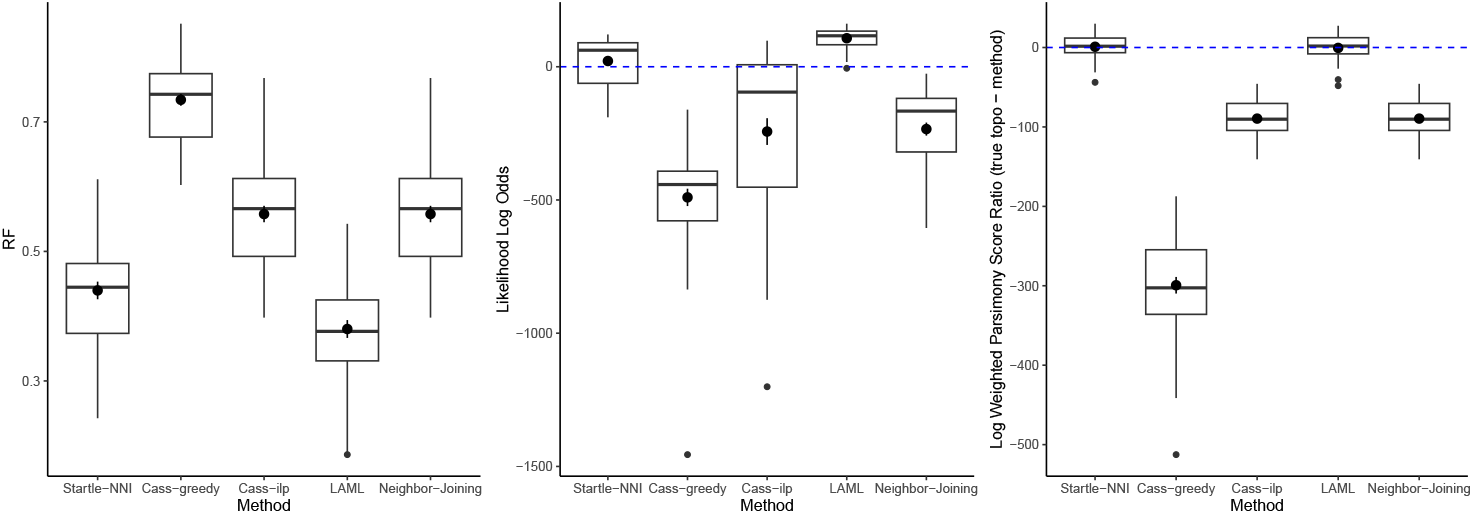
Comparison of the benchmarked methods based on two metrics: estimated log likelihood (calculated by LAML) to the true tree likelihood on simulated data (left); and weighted parsimony score (right).

#### S2.6.1 Branch Length Results

**Figure S4:**
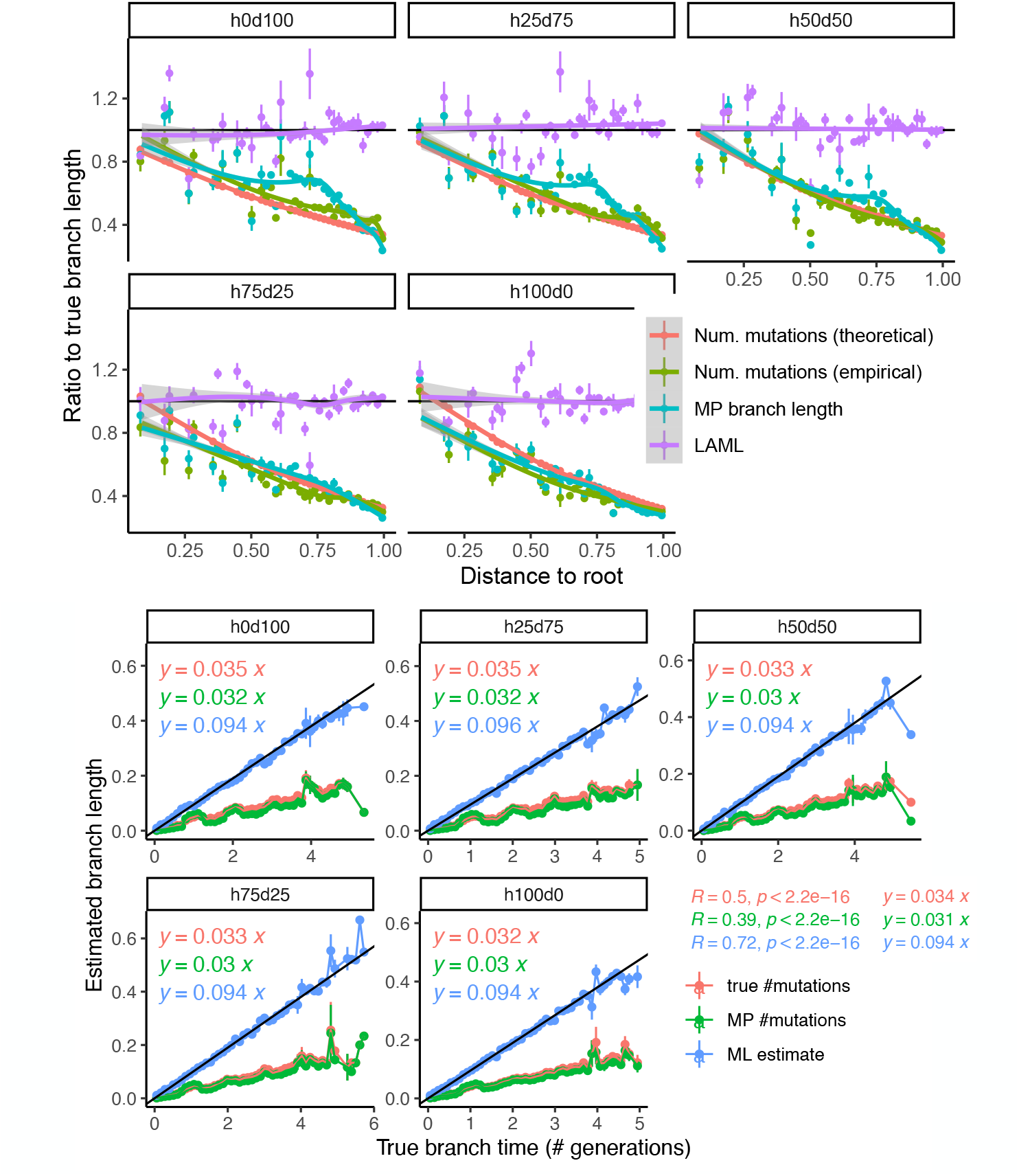
Branch length estimation analysis on true topologies. Top: The the ratio of estimated to true branch length versus distance to root. LAML’s estimate (i.e. ML estimate, shown in blue) is compared to two alternatives: true # mutations and MP # mutations, which are the true number of mutations and the estimate by maximum parsimony respectively. The x-axis is discretized into 50 bins and the estimated branch length by each method is averaged for each bin, shown with error bar. The black line shows the perfect estimated/true = 1. Bottom: regression of the estimated versus true branch length. The x-axis is discretized into 50 bins and the estimated branch length by each method is averaged for each bin, shown with error bar. The black line shows the ideal fit using the true known mutation rate *λ* = 0.095. Correlation coefficient (*R*), p-value (*p*), and the equation of the linear regression are shown for each method.

#### S2.6.2 Runtime Results

Fig S5 illustrates the breakdown of the EM algorithm into the E-step and two cases of the M-step. This is on the model condition with zero heritable and 100 dropout (h0d100). We ran on varying number of cells (up to 5000 subsampled cells). The M-step non-convex case does not have a value for the largest case as those jobs did not complete in the 7 days of runtime provided.

**Figure S5:**
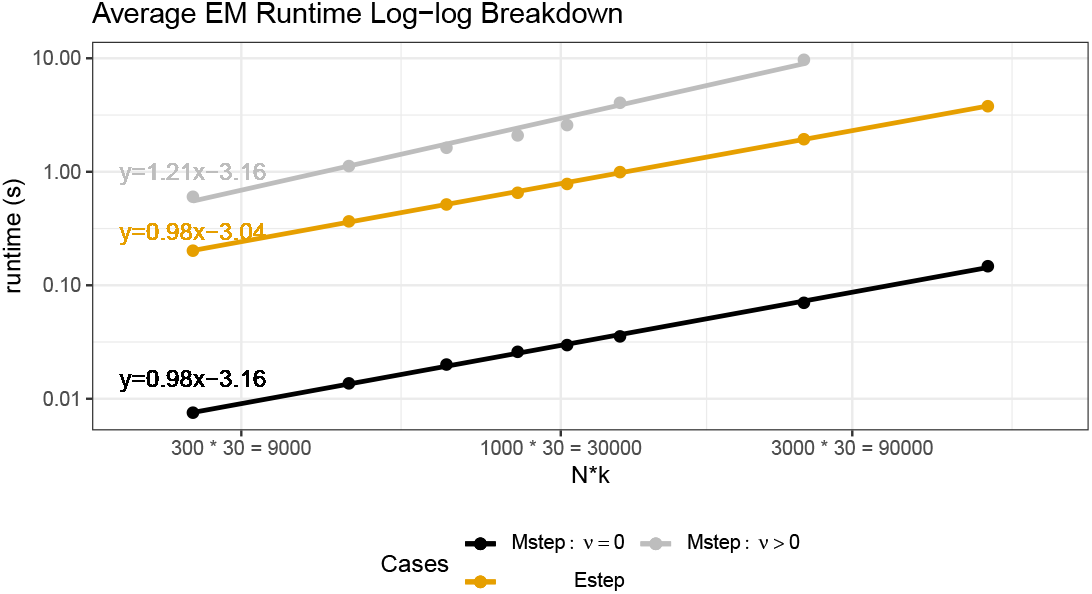
Runtime of EM algorithm, shown for E-step and M-step with the non-convex heritable and convex non-heritable cases (log-log plot). The slope of the fitted lines are all around 1, indicating that empirical runtime of these steps are all close linear.

We analyze the runtime efficiency of the EM algorithm first, then provide an exploration of the total topology search runtime. Fig. S6 illustrates how runtime and EM iterations changes as the number of cells increase.

**Figure S6:**
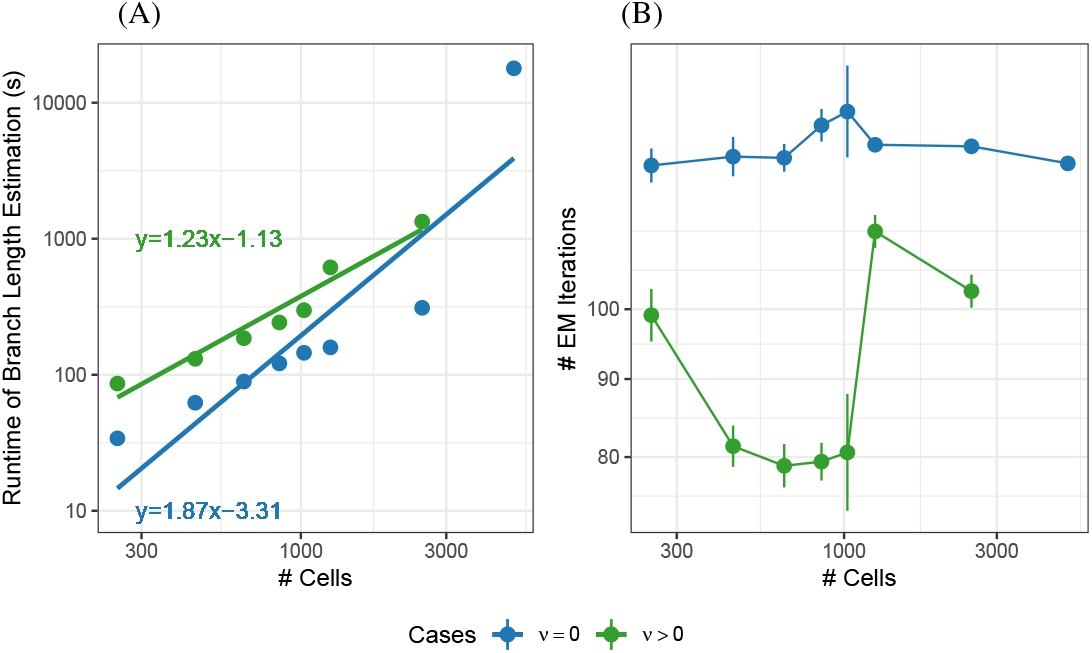
Empirical runtime (A) and number of EM iterations (B) of LAML on simulated data with varying number of cells (up to 5000 cells).

We provide a runtime comparison of topology search algorithms (Cassiopeia-Greedy, LAML, Startle-NNI and Cassiopeia-ILP). We run on simulated data sub-sampled to 250 cells, across all model conditions but only on replicates (≈ 40) for which Cass-ILP successfully completed.

**Figure S7:**
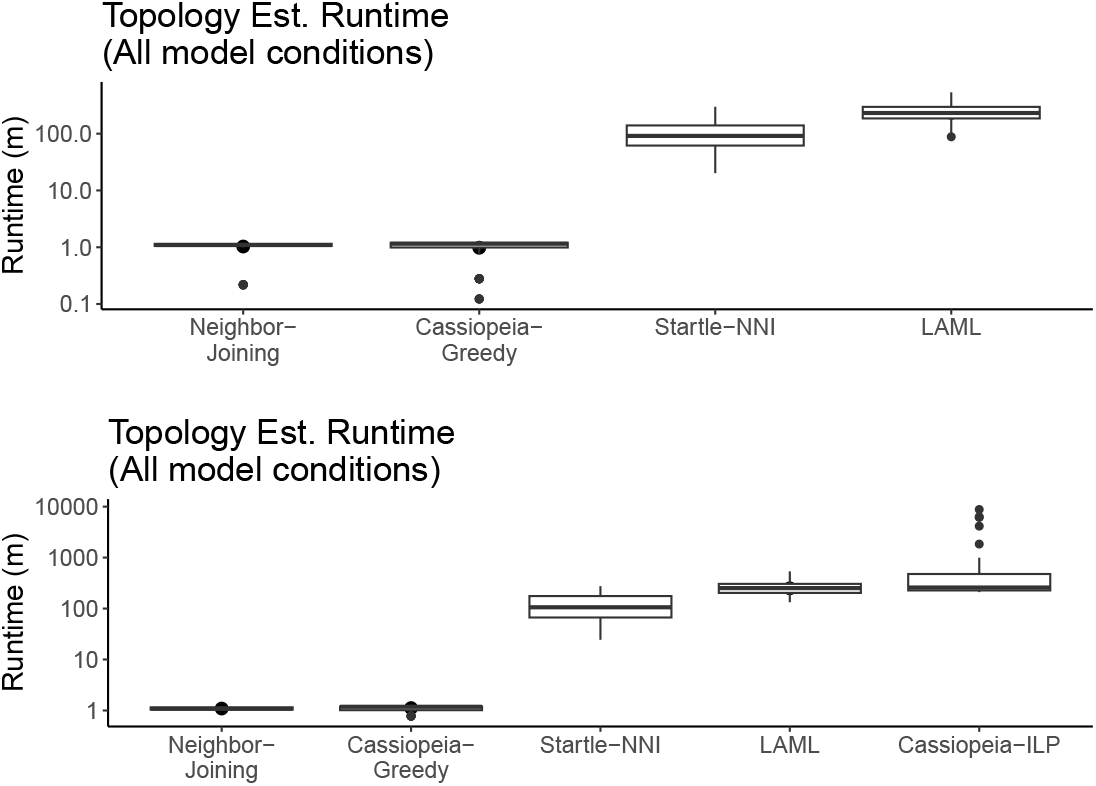
Average runtime comparison for topology estimation on simulated data. Runtime is shown in minutes in log scale.

### S2.7 Supplementary Results for KP-Tracer

**Table S4:** Migration cost of the trees inferred by Cass-hybrid, Startle-NNI, and LAML on KP-tracer sample 3724 NT T1 All.

|  | Migration Types | LAML | Cass-hybrid | Startle-NNI |
| --- | --- | --- | --- | --- |
| De-duplicated<br>(1,461 cells) | All migrations | **99** | 136 | 119 |
|  | Reseeding | **42** | 68 | 56 |
| Full Tree<br>(21,108 cells) | All migrations | **1568** | 1600 | 1783 |
|  | Reseeding | **191** | 219 | 203 |

#### S2.7.1 Migration Metrics Table

From Table S5, we can observe that the LAML tree topology always achieves a lower migration cost than the Cassiopeia-Hybrid tree does.

**Table S5:** Metrics computed on the *MG*_*SC*_ migration graph with spatial clustering of all observed cells.

| Metrics | LAML ($T_{LAML}$) | Cassiopeia-Hybrid ($T_{CassH}$) |
| --- | --- | --- |
| Total Cost | 836 | 959 |
| Induced Anatomical Cost | 125 | 159 |
| Spatially-aware Migration Cost (Primary Tumor) | 730 | 828 |
| Constrained Parsimony | 935 | 1095 |

#### S2.7.2 Metastasis Analysis

With the knowledge that this experiment ran for 6 months, we scale our ultrametric tree, which had root-to-tip distance of 2.47 to be 6 with a scale factor of 2.425. To produce the bottom half of Figure 4(B), we begin with our tree with inferred anatomical labels on the internal nodes as described in the main text [14] and in Supplementary Section S2.1.2.

**Figure S8:**
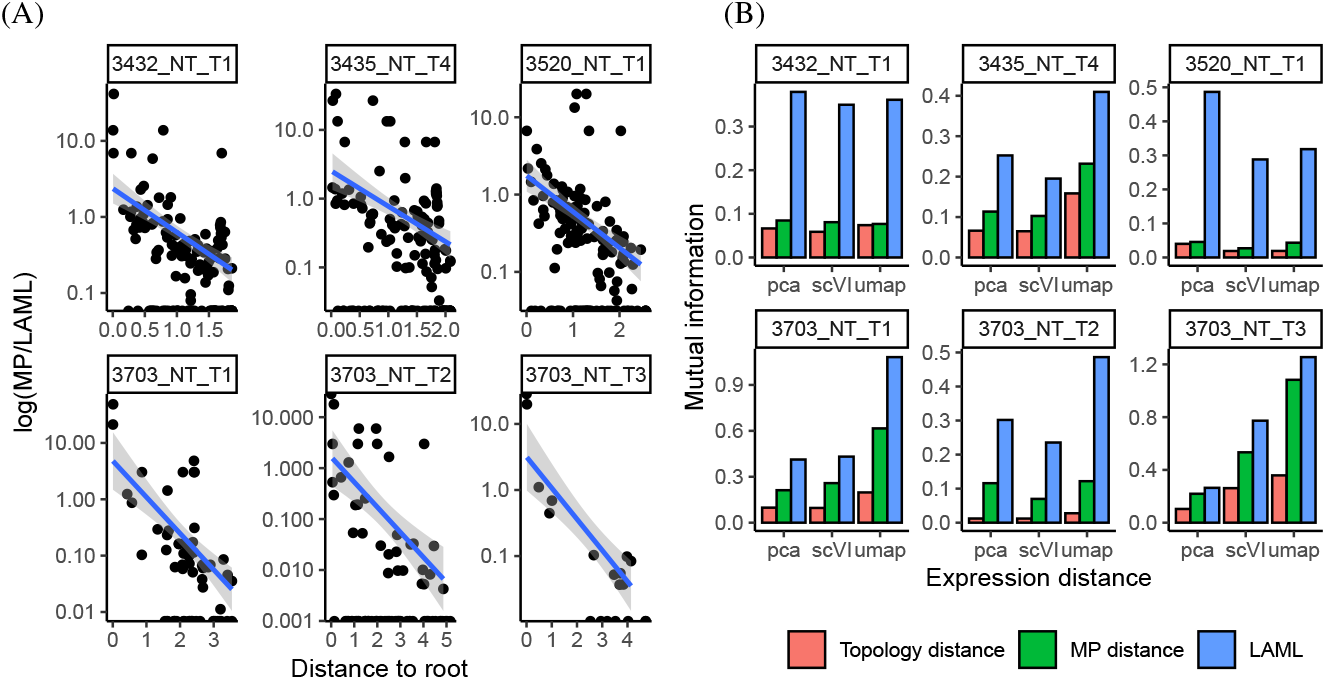
Branch length estimation by LAML (dML) and Maximum Parsimony (dMP) on the 6 selected samples of KP-tracer. (A) The ratio of each cell’s branch length (i.e. the branch above the node) estimated by maximum parsimony (dMP) and maximum likelihood (dML) versus its distance to the root. The y-axis is shown in log-scale. (B) Mutual information of pairwise gene expression and phylogenetic distances. Gene expression is represented either by PCA, scVI, or UMAP. For these 6 samples, we use the published tree topology (estimated by Cassiopeia) and estimate branch lengths by LAML (ML distance), maximum parsimony (MP distance), or solely use the tree topology (Topology distance).

We annotate each edge *e* = (*u, v*) based on the anatomical labels of the parent node *u* and child node *v*. We use edge, branch and lineage interchangeably in this section.

Notably, we are interested in transition edges, where one of the nodes is labeled with the primary tumor anatomical site, and the other node is labeled with one of the other 4 anatomical sites. Thus, we define four edge types: non-transition edge, primary to non-primary (metastasis), non-primary to primary (reseeding), and non-primary *i* to non-primary *j* (where *i ≠ j*).

Next, we establish 2, 470 intervals ranging from 0.0 to 2.47 with a step size of 0.001. In each interval, we find the branches which overlap with this interval. Thus, we can compute the number of each branch type which overlap with this interval. Since the tree grows to include more cells in each cell generation, we normalize the counts of each edge type by the total number of branches overlapping this interval.

We can do an additional analysis to compute the expected number of unedited sites left at any given time point. We use the following equation: exp( −*d/c*) * 9, where *d* is the time-scaled distance to root, and *c* is our scaling factor. It is useful to consider the expected number of unedited sites left at various time points (i.e. at the 3 month mark in the main paper), to give a sense of whether there is still phylogenetic signal left.

We also analyze the length of any branches which lie entirely within a single epoch in Figure S9. Note that the branches in the Late Metastasis epoch are much longer as expected due to the loss of phylogenetic signal and subsequent low tree resolution. The branch lengths observed in the the Primary Growth epoch are shorter, suggesting the phylogenetic signal is capturing cell divisions. Interestingly, the Metastasis epoch has the shortest branches *<* 1.0 month, perhaps suggesting a general increase in cell division rates. The Metastasis Burst window within the Metastasis epoch has even fewer outlier branch lengths.

### S2.8 Supplementary Results for intMEMOIR

Comparison with TiDeTree on all the intMEMOIR samples. Note that the TiDeTree topologies shown here are the maximum credibility clade trees obtained from the TiDeTree authors, which show trees jointly estimated (sharing parameters across all samples). We ran Cassiopeia-Hybrid with a cut-off of 100 cells, so that we effectively report results for Cassiopeia-ILP on all intMEMOIR samples.

**Figure S9:**
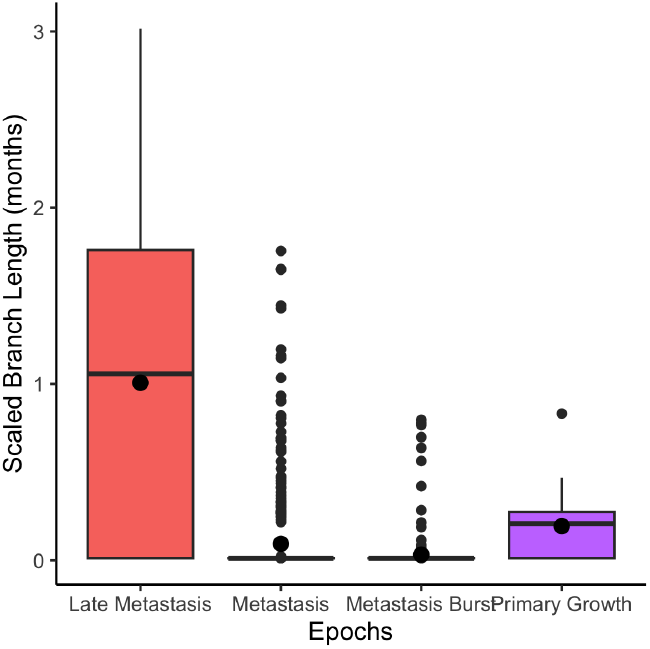
Length of branches lying entirely within each epoch

**Figure S10:**
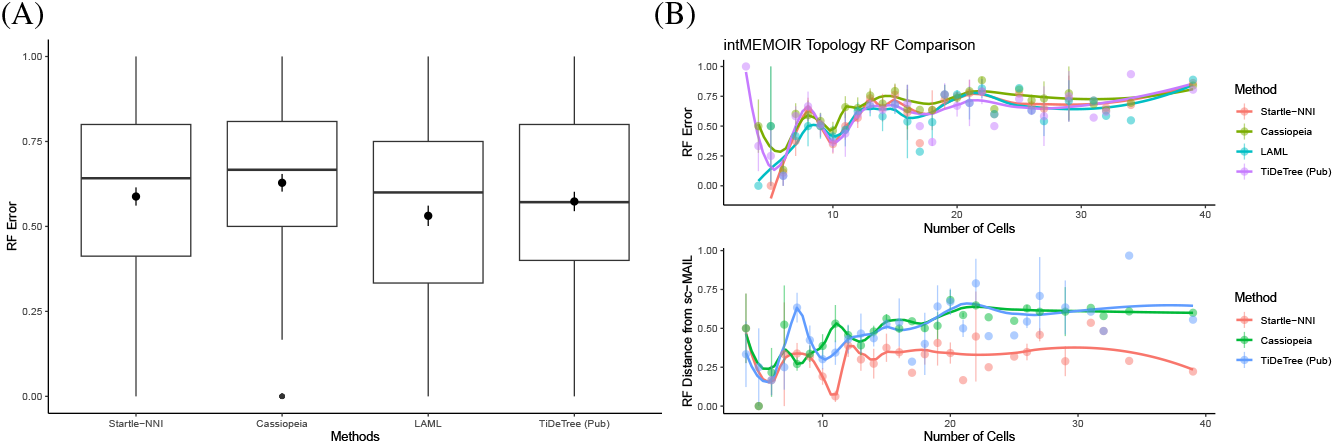
(A) Topology accuracy (RF) comparison between the true tree and trees estimated using LAML and the benchmarked methods, on all intMEMOIR samples. (B) RF distance plotted against the varying number of cells. The top plot shows RF distance from the true tree and the bottom plot shows RF distance from the LAML tree.

Runtime comparison on the intMEMOIR data (see Fig S11).

**Figure S11:**
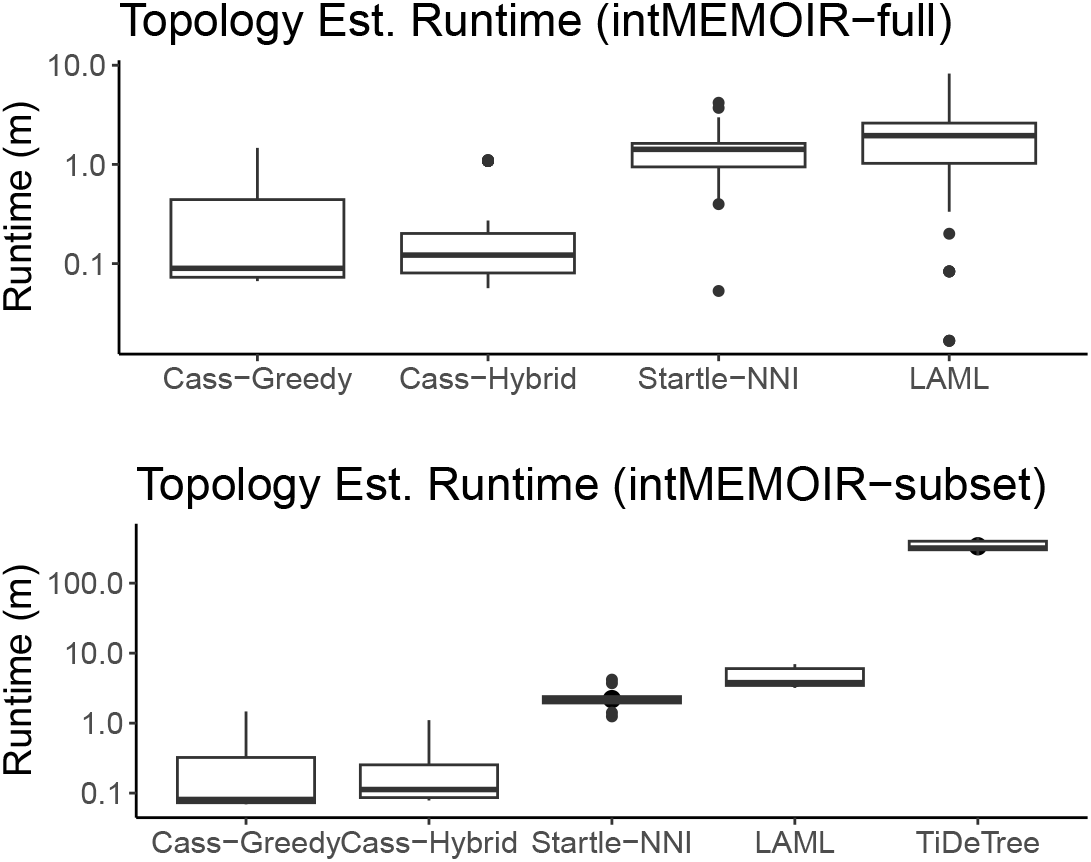
Topology estimation runtime comparison between LAML and TiDeTree on intMEMOIR. Note that the runtime of each methods’ starting tree has also been added to that method’s runtime.

### S2.9 Distance-Based Exploration

**Figure S12:**
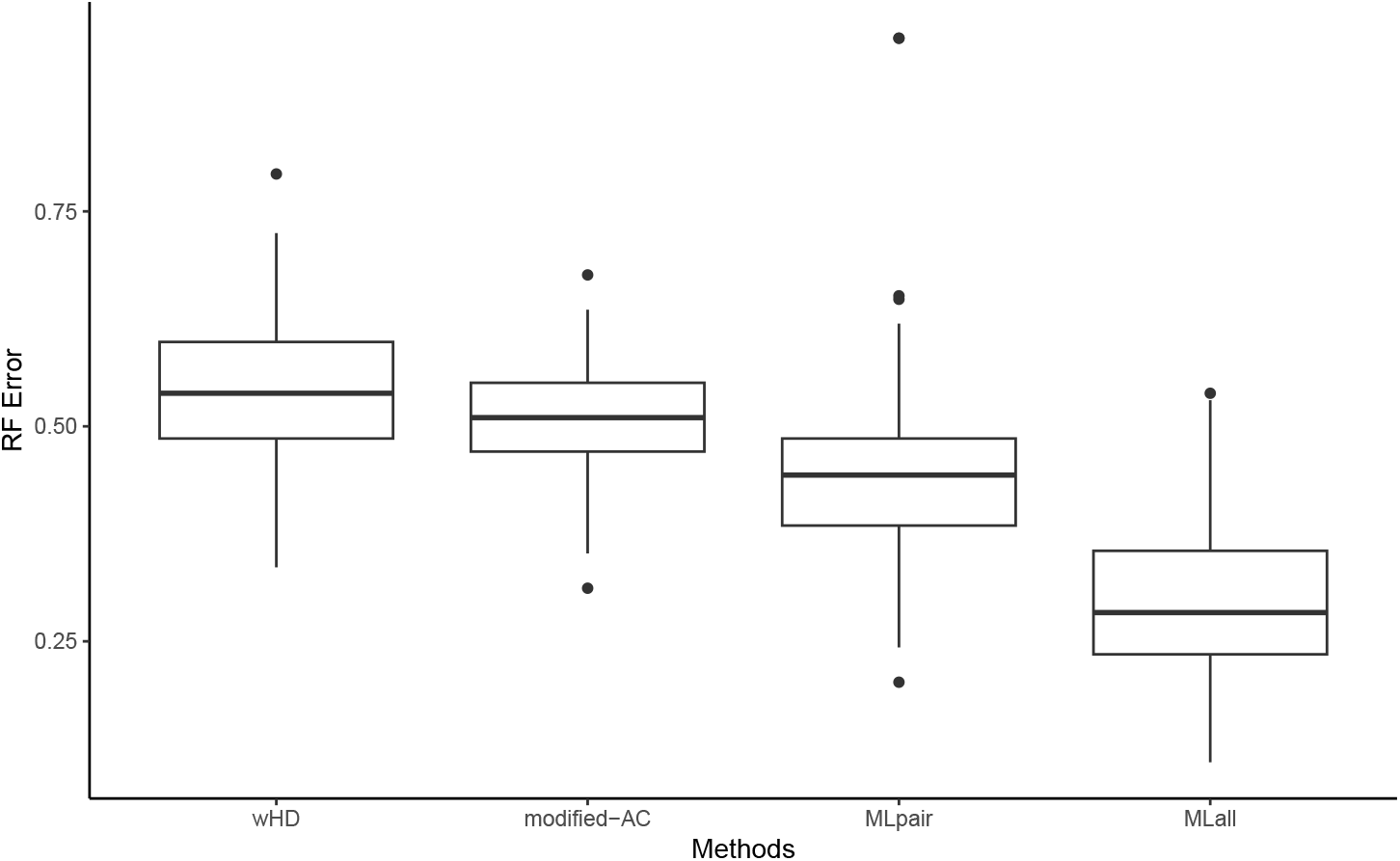
Comparison of different approaches to distance estimation. Two pipelines: (1) From the left, the first three pipelines first estimate rooted pairwise distances, before tree estimation using NJ. (2) MLall (LAML) finds the maximum likelihood distance matrix and tree topology over all pairs of leaves at once.

Figure S12 illustrates how distances estimated under this statistical model improve tree estimation. wHD, which does not take the missing state into consideration, produces trees with the highest RF error across all methods. This can be contrasted with the mAC, which does consider the missing state. The MLpair approach, which explicitly models the two types of missing data for each pair of cells, further improves the RF error. As expected, the MLall approach, which is able to optimize distances between sequences over all pairs of sequences, achieves the lowest RF error. This indicates the importance of estimating distances under a statistical model explicitly modeling the two types of missing data.

### S2.10 Review of Lineage Tracing Technologies

Dynamic lineage tracing technologies begin by engineering one or more progenitor cells with artificial recording sites, then accumulating heritable insertions or deletions (“indels”) [46] over cell generations. We classify dynamic lineage tracing technologies into 3 generations of technology. In the first generation of dynamic lineage tracing (GESTALT[28] and MEMOIR [17]), the recordable time is low because of limited number of target sites (no more than 10) and edit states (limited to 2: inversion and deletion). In the second generation (scGESTALT [35], LINNAEUS [44], ScarTrace [2], and intMEMOIR[10]), other information such as gene expression and spatial information, are recorded in addition to CRIPSR-Cas9 edit, enhancing tree inference [35]. However, the recordable tree depth remains low (only 6 cell generations) [28, 35], and there is a high error rate as edits are often removed or overwritten [44, 46]. In the third generation, novel dynamic lineage tracing technologies, such as CARLIN [5], iTracer [23], Chan et al. [7], Yang et al. [52] and TypeWriter [9], substantially improve recordable tree depth. We can further classify these technologies into two groups. The first group comprises technologies that induce editing at distinct time points. The second group comprises TypeWriter [9] and Chan et al. [7], the technologies focusing on increasing the number of cell generations by optimizing the number of editable sites and states. According to TypeWriter, the recordable tree depth can scale to 20 generations. The technology developed by Chan et al. is later demonstrated in [52], where number of target sites and edit states scale up to hundreds, and the recordable tree depth scales to 15 generations. Notably, the largest sample has 21,108 cells spanning 5-6 months of cell growth.

We should note that although the number of editable target sites affects the number of cell generations one can record over, that the quality of target sites affect how many of them turn out to be informative. Note also the distinction between continuous editing and inducible editing approaches: the depth of a given reconstructed cell lineage tree may span different numbers of cell generations and developmental time. **PMM is applicable to *all* of these lineage tracing technologies**.

**Table S6:** Summarizing key properties of the different lineage tracing technologies.

| Lineage Tracing Technology | Lineage Tree Depth | Time Recorded Over | # Target Sites | Approach | Size of Edit Library | Edits Control |
| --- | --- | --- | --- | --- | --- | --- |
| ScarTrace | 2 | 10 hours | 8 | barcode | uncontrolled | Cas9 injection |
| LINNAEUS | 6 | 10 hours | 16-32 | CFP region | uncontrolled | Cas9 injection |
| CARLIN | 3 | / | 10 | barcode | uncontrolled | dox inducible |
| iTracer | 5 | 100 hours | 4 (Fig 6., Ext. Fig 10.) | barcode | uncontrolled | dox inducible |
| scGESTALT | 5 | / | 9 | barcode | uncontrolled | heat shock inducible |
| Chan et al. | 12 | 6 days | 9-45 | barcode | uncontrolled | continuous |
| Yang et al. | 15 | 6 months | 10-30 | barcode | uncontrolled | continuous |
| intMEMOIR | 4 | 36 hours | no target sites<br>10 transgenes | barcode<br>not CRISPR edits | uncontrolled | heat shock inducible |
| Typewriter | 20 | 25 days | 48 “tapes” | prime editing | by design: prototyped up to 16 | dox inducible |

### S2.11 Review of Other Computational Approaches

Computational methods for lineage tracing can be broadly categorized into three groups: maximum parsimony (MP), distance-based, and probabilistic modeling, with some methods exhibiting hybrid characteristics. Maximum parsimony methods [25, 39] rely on the heuristic criterion of maximum parsimony (MP). Cassiopeia [25], a highly regarded method in the DREAM challenge [20], infers the MP tree using a modified Camin-Sokal (C-S) model to leverage the non-modifiability property of CRISPR/Cas9 edits. Sashittal et al. [39] later refer to this evolutionary model as the “star homoplasy model” and develop Startle [39] to find the MP tree. However, due to the use of the heuristic MP criterion, neither Cassiopeia or Startle can adequately capture the stochastic nature of the CRISPR/Cas9 process. The second category encompasses distance-based methods, such as neighbor joining and triplet-based approaches. A critical challenge in these methods lies in computing pairwise distances, which are essential for generating the distance matrix (for neighbor joining) or triplet topologies (for triplet-based methods). Recent research [**?**] has shown that accurate computation of distances can be achieved with theoretical guarantees, but only when a probabilistic model is assumed. The last category comprises methods that formulate lineage tracing as a statistical inference problem. Bayesian [40] and maximum likelihood [16, 26, 52] methods fall within this category. Many of these methods model a specific set of lineage tracing technologies and do not generalize well to other technologies. For example, GAPML [16] introduced a probabilistic model tailored for the scGESTALT technology, focusing on single-barcode evolution. LiNTIMaT [52] can only be used for the second generation of lineage tracing technologies, where there are only two edit states, and requires expression data as input. TiDeTree [40] is designed more generally for barcode-based technologies, but their model assumes sites are independent and *identically distributed*, limited its application to the handful of technologies having all target sites shared a same set of edit states.

## Footnotes

1 Following the convention in phylogenetics, we use *target site* and *site* interchangeably.

2 Throughout this paper, we use *edge* and *branch* interchangeably.

